# Predation drives specialized host plant associations in preadapted milkweed bugs (Heteroptera: Lygaeinae)

**DOI:** 10.1101/2020.06.16.150730

**Authors:** Georg Petschenka, Rayko Halitschke, Anna Roth, Sabrina Stiehler, Linda Tenbusch, Tobias Züst, Christoph Hartwig, Juan Francisco Moreno Gámez, Robert Trusch, Jürgen Deckert, Kateřina Chalušová, Andreas Vilcinskas, Alice Exnerová

## Abstract

Host plant specialization across herbivorous insects varies dramatically, but the underlying evolutionary mechanisms are little-known. The milkweed bugs (Heteroptera: Lygaeinae) are ancestrally associated with plants of the Apocynaceae from which they commonly sequester cardiac glycosides for defense, facilitated by resistant Na^+^/K^+^-ATPases and adaptations for transport, storage and discharge of toxins. Here, we show that three Lygaeinae species independently colonized four novel non-apocynaceous hosts, convergently producing cardiac glycosides. A fourth species shifted to a new source of toxins by tolerating and sequestering alkaloids from meadow saffron (*Colchicum autumnale*, Colchicaceae). Across three species tested, feeding on seeds containing toxins did not improve growth, but sequestration mediated protection against predatory lacewing larvae and birds. We conclude that physiological preadaptations and convergent phytochemistry facilitated novel specialized host associations. Therefore, selection by predators on sequestration of defenses, rather than the exploitation of novel dietary resources, can lead to obligatory specialized host associations in generalist insects.

## Introduction

Herbivorous insects show tremendous variation with regard to dietary specialization. While it is a long-standing assumption that phytochemicals may restrict and direct the evolution of host plant use^1^, the explicit role of phytochemicals as drivers of host plant associations has been revealed in only a few systems^2–5^. Proposed mechanisms of how plant secondary compounds could mediate insect-plant interactions include physiological trade-offs in the efficiency of host plant use between generalists and specialists^1,6–9^. Alternatively, it has been shown that novel host plant associations can create enemy-free spaces for herbivores^10,11^ either by providing defense^11^ or refuge from natural enemies^10^. However, even though it is widely recognized that many insects not only use plants as a dietary resource but also sequester (i.e. absorb and store) plant toxins to defend themselves against predators^12–14^, the extent to which sequestration could drive the evolution of insect-host plant associations has rarely been addressed^4,15^.

While it has been hypothesized that dietary specialization and sequestration of plant toxins can lead to an evolutionary dead end^4,16^, there is evidence that ecological specialization does not necessarily prevent host range expansion^4^. Nevertheless, sequestration and dietary specialization seem to be evolutionarily linked^13,17–20^, and predators driving the occupation of enemy-free-spaces are typically considered to select for specialization^9,21^. Recent research indicated that sequestration requires different resistance traits than are required to merely cope with dietary toxins^22^, suggesting that selection by predators or parasitoids (i.e. the third trophic level) opens a second arena for coevolutionary escalation^13,22^. Consequently, a rigorous analysis of coevolution between plants and specialized insects requires the integration of adaptations underlying bitrophic interactions with adaptations driven by higher trophic levels^13,22,23^.

Here, we used milkweed bugs (Heteroptera: Lygaeinae) as a model system to test hypotheses about the evolutionary drivers leading to specialized associations with particular plant species. The Lygaeinae comprise about 600 primarily seed-eating species that are well known for their predilection of plants in the Apocynaceae worldwide^24–27^. Milkweed bugs typically exhibit a red-and-black aposematic coloration and, in addition to defensive scent glands typical for Heteroptera^28^, several species have been shown to acquire defenses against predators from their host plants^29–32^. Upon attack, many milkweed bug species release sequestered toxins in a defensive secretion from a specialized storage compartment of the integument (the dorsolateral space)^33,34^. The large milkweed bug (*Oncopeltus fasciatus* (Dallas, 1852)) in particular has been studied in detail with regard to sequestration of cardiac glycosides, which it derives from seeds of milkweeds in the Apocynaceae genus *Asclepias*^35^.

Cardiac glycosides are important defense metabolites of plants in the Apocynaceae, and evolved convergently in at least 11 additional botanical families^36^. Both compound subtypes, the cardenolides and the bufadienolides, are specific inhibitors of the ubiquitous animal enzyme Na^+^/K^+^-ATPase. Specialized insects from at least six taxonomic orders, including several lygaeine species^37–40^, tolerate cardenolides by expressing Na^+^/K^+^-ATPases with several amino acid substitutions that mediate a high degree of cardenolide resistance in vitro (target site insensitivity)^40,41^. Duplication of the gene coding for the α-subunit of Na^+^/K^+^-ATPases in the lygaeine bugs *O. fasciatus* and *Lygaeus kalmii* Stal, 1874 resulted in three different copies with up to four amino acid substitutions in regions of the protein critical for cardiac glycoside binding^38,39,42^, rendering the enzyme resistant to cardiac glycosides^42^. In addition to resistance, sequestration requires accumulation of toxins from the dietary resource, and milkweed bugs possess an as-of-yet unidentified mechanism for the transport of toxins across the gut epithelium. In summary, milkweed bugs possess a suite of traits related to sequestration and defense that includes aposematic coloration, resistant Na^+^/K^+^-ATPases, and mechanisms for accumulation, storage, and release of toxins. This suite of traits may also function as a physiological preadaptation facilitating the sequestration of novel toxin compounds. For example, the milkweed bug *Neacoryphus bicrucis* (Say, 1825) sequesters pyrrolizidine alkaloids^31^, a class of compounds unrelated to cardiac glycosides.

Sequestration of cardiac glycosides by lygaeine bugs was initially described for *O. fasciatus* and *L. kalmii* feeding on *Asclepias* species^43^, as well as for *Caenocoris nerii* (Germar, 1847) and *Spilostethus pandurus* Scopoli, 1763 on oleander (*Nerium oleander*), both belonging to the Apocynaceae^44^. A broad survey based on dried museum specimens demonstrated the presence of cardiac glycosides in many genera of Lygaeinae^24^, suggesting that sequestration of cardiac glycosides is a common trait of milkweed bugs. Furthermore, an evolutionary analysis revealed that sequestration of cardiac glycosides, target site insensitivity of Na^+^/K^+^-ATPase, as well as an association with apocynaceous plants are likely ancestral traits of the group^40^. Nevertheless, despite being specialized to cardiac glycosides, several milkweed bug species are dietary generalists and feed on seeds from a great variety of plant families. The palaearctic species *Lygaeus equestris* (Linnaeus, 1758) for example, was observed feeding on more than 60 plant species from roughly 20 families^45^. Even though dietary breadth can vary among species and larval instars to some extent^26,45^, this large number of potential host plants is in stark contrast to the narrow set of species that produce cardiac glycosides for sequestration. The extreme prevalence of cardiac glycoside sequestration in milkweed bugs thus suggests an important role of selective pressure exerted by higher trophic levels in shaping milkweed bug-host plant associations.

Remarkably, several palaearctic species of Lygaeinae are regularly found on plants which are phylogenetically disparate from the Apocynaceae, but which convergently produce cardiac glycosides. *Horvathiolus superbus* (Pollich, 1779) was observed feeding on *Digitalis purpurea* (Plantaginaceae) and seems to depend on this plant at least in parts of its distributional range^26,46^. In addition, we found this species using the cardenolide-producing *Erysimum crepidifolium* (Brassicaceae) as a host plant in a *Digitalis-free* habitat. Early instars of *L. equestris* larvae in Sweden feed exclusively on *Vincetoxicum hirundinaria* (a cardenolide-free Apocynaceae) and the cardenolide producing Ranunculaceae *Adonis vernalis*^45,47^ which is also an important host plant elsewhere^26^. The generalist milkweed bug *S. pandurus* was recorded on *Urginea maritima* (Asparagaceae)^48^ which produces cardiac glycosides of the bufadienolide-type^49^.

Surprisingly, a closely related species, *Spilostethus saxatilis* (Scopoli, 1763), which uses a great variety of host plants^26,48^, is not known to visit cardiac glycoside producing plants except for *Asclepias syriaca*^50^ that is not native in its distributional range. However, we and others^26,51^ have often observed this species on flowers and fruits of meadow saffron (*Colchicum autumnale*, Colchicaceae) which is highly toxic due to the production of alkaloids such as colchicine. Colchicum alkaloids inhibit polymerization of tubulin^52^, thus showing a mode of action that is different from cardiac glycosides. Nonetheless, based on its evolutionary history, *S. saxatilis* may be preadapted regarding some traits of the ‘sequestration suite’ including aposematic coloration, storage and release of toxins, and a putative mechanism for transport. In a scenario of sequestration-driven host associations and selection by higher trophic levels, colonization of new sources of potent toxins for sequestration could represent a mode of coevolutionary escalation.

Using these insect-plant interactions as a model system, we tested if three species of milkweed bugs sequester cardenolides from four independently colonized, cardiac glycoside-containing plants in the Asparagaceae, Brassicaceae, Plantaginaceae, and Ranunculaceae, and furthermore tested if *S. saxatilis* sequesters alkaloids from *C. autumnale*. After confirming the tolerance and sequestration of colchicum alkaloids by *S. saxatilis*, we evaluated the degree of association between *S. saxatilis* and *C. autumnale* by screening 30 *S. saxatilis* museum specimens from 11 countries for the presence of both colchicum alkaloids and cardenolides. We also assessed if there is an ecological trade-off between sequestration of colchicum alkaloids and cardiac glycosides in this species. Next, we evaluated the importance of cardiac glycoside- or alkaloid-bearing seeds as a dietary resource for milkweed bugs by quantifying larval growth of *H. superbus*, *L. equestris*, and *S. saxatilis* fed with different combinations of toxic- and non-toxic seeds. We then contrasted the effects on growth with a quantification of defensive benefits gained by consumption and sequestration of cardiac glycosides and colchicum alkaloids in *H. superbus, L. equestris*, *S. pandurus*, and *S. saxatilis* against insect (lacewing larvae, *Chrysoperla carnea*) and avian predators (great tits, *Parus major*). Finally, we tracked the evolution of colchicine resistance in a phylogenetic framework using cardenolide and colchicine injection assays to mimic sequestration.

We suggest that sequestration of plant toxins as a defense against predators mediates specialized associations of milkweed bugs with specific host plant species. The preadaptations for sequestration of cardiac glycosides and their convergent occurrence within distantly related plants most likely facilitated new host associations. Furthermore, a subset of these preadaptations may have facilitated the shift to new host plants with functionally distinct but highly potent toxins. We thus demonstrate that species which are dietary generalists under a bitrophic perspective may nonetheless be highly specialized on plants that provide defenses against the third trophic level.

## Material and Methods

### Field-sampling of milkweed bugs for chemical analysis

In order to assess sequestration of toxins under natural conditions, adult milkweed bugs were collected in the field from habitats with natural stands of their toxic host plants (Figure 1, Supplemental Figure 6). Field work in protected areas was permitted by the responsible agencies (see acknowledgments). Notes on natural history observations are given in the electronic supplementary material. *S. saxatilis* were collected in Nüstenbach (August 15^th^, 2015) and Berghausen (September 18^th^, 2015), Baden-Württemberg, Germany, either from *C. autumnale* (Colchicaceae) or feeding on other plants. *H. superbus* was collected on August 15^th^ and 16^th^, 2015 in a *Digitalis purpurea* (Plantaginaceae) habitat close to Eberbach, Baden-Württemberg, Germany. We collected the same species feeding on *Erysimum crepidifolium* (Brassicaceae) in a *Digitalis-free* habitat (Schloßböckelheim, Rheinland-Pfalz, Germany) on June 12^th^ and 13^th^ 2018. *L. equestris* was collected in the nature reserve ‘Oderhänge Mallnow’ north of Lebus, Brandenburg, Germany on April 19^th^, 2016 during the blooming of *Adonis vernalis* (Ranunculaceae). Lastly, *S. pandurus* was collected from infructescences of *Urginea maritima* (Asparagaceae) in ‘Parque Natural Sierra de Aracena y Picos de Aroche’, Aracena, Spain, in early November 2016. After bringing the bugs to the lab, they were maintained on sunflower seeds and water provided in Eppendorf tubes plugged with cotton under ambient conditions to purge their guts from remaining toxins. After 14 days (≥ 12 days for *S. pandurus*) bugs were frozen at −80°C and freeze-dried for chemical analysis as described below.

**Figure 1.**
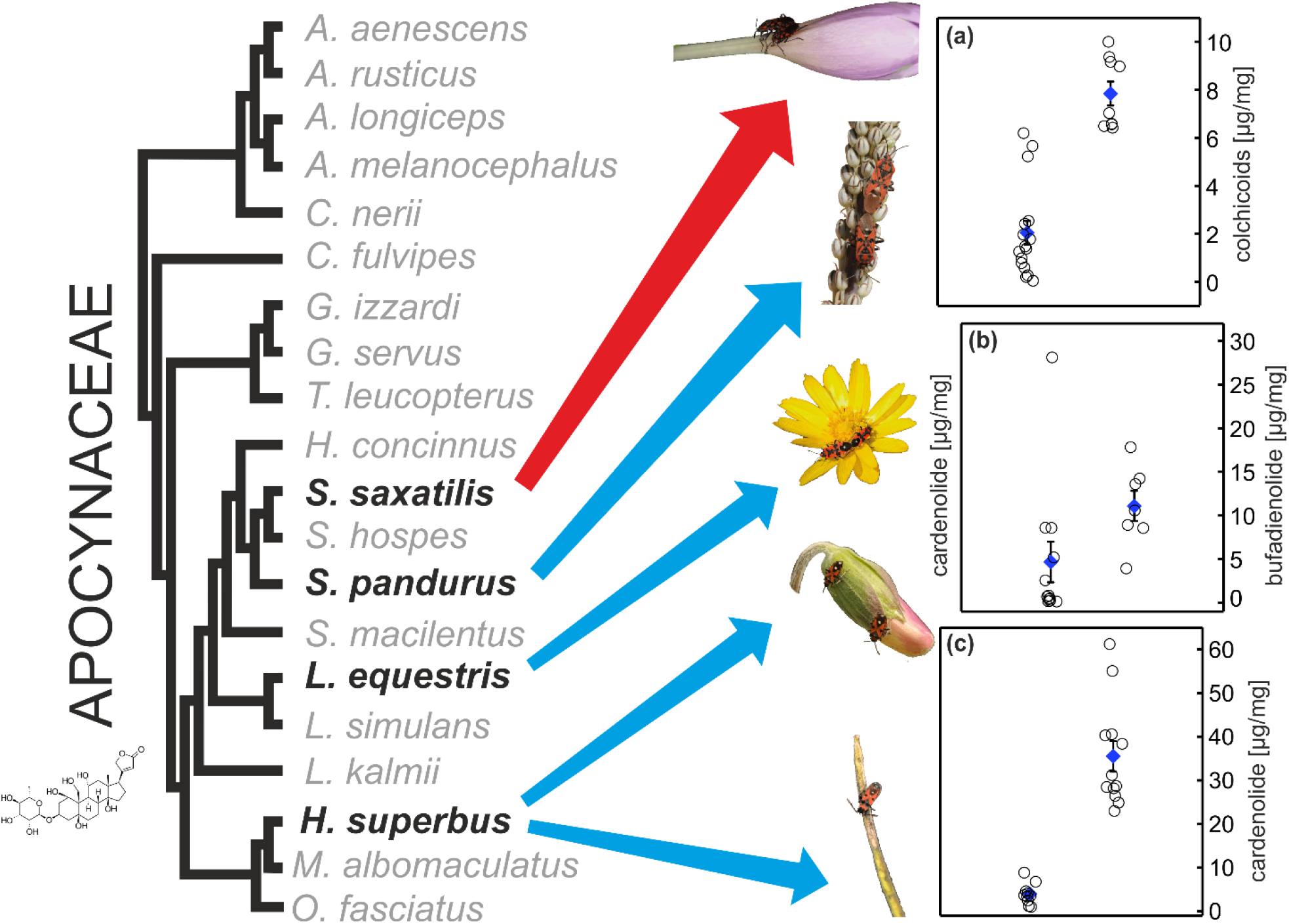
Association of milkweed bugs with toxic plants and sequestration of toxic plant compounds in field-collected specimens. Left and center: phylogenetic relationships of milkweed bugs^40^ and host plants of focal species. The red arrow indicates the association of *S. saxatilis* with the colchicum alkaloid producing *C. autumnale* (Colchicaceae), blue arrows indicate associations with plants containing cardiac glycosides. From top to bottom: *S. saxatilis* on *C. autumnale*, *S. pandurus* on *U. maritima* (Asparagaceae), *L. equestris* on *A. vernalis* (Ranunculaceae), *H. superbus* on *D. purpurea* (Plantaginaceae), and *H. superbus* on *E. crepidifolium* (Brassicaceae). Right: Concentrations of plant toxins sequestered by milkweed bugs collected in the field. (a): Colchicum alkaloids in *S. saxatilis* from two different populations (left: Nüstenbach, n = 11, right: Berghausen, n = 9). (b): Cardenolides in *L. equestris* from *A. vernalis* (left, n = 12) and bufadienolides in *S. pandurus* from *U. maritima* (right, n = 7). Scale for cardenolides is identical with scale for bufadienolides. (c): Cardenolides in *H. superbus* from *D. purpurea* (left, n = 10) and in *H. superbus* from *E. crepidifolium* (n = 12). Diamonds represent means ± SE, circles represent jittered raw data.

### Origin and maintenance of bugs used for experiments

Specimens of *L. equestris, H. superbus*, and *S. pandurus* used for seed mixture feeding, predation, and injection experiments were obtained from laboratory colonies maintained for several generations at the time of the experiments. Specimens for founding colonies were collected at the locations described above (*H. superbus* from Eberbach), except for *S. pandurus* used in injection and predation assays that were from a lab-strain originating from Portugal collected in 2016. *Oncopeltus fasciatus* used as controls for injection assays were from a long-term laboratory colony (origin United States), obtained from the University of Hamburg in 2015. Bugs were raised on husked sunflower seeds and supplied with water (see above) as well as pieces of cotton for oviposition. Colonies were maintained in environmental chambers at 28°C and 60% humidity at a light: dark cycle of 16 h: 8 h. Specimens of *S. saxatilis* were raised from eggs obtained from adults collected in the field (Berghausen) that were fed with sunflower seeds under ambient conditions. *Pyrrhocoris apterus* (Linnaeus, 1758) used as a non-adapted outgroup for injection assays were collected in the field (Giessen, Germany) and used directly or after one day of maintenance in the laboratory on linden seeds.

### Seed mixture experiments

To assess if inclusion of *Digitalis*-, *Adonis*-, or *Colchicum*-seeds (‘toxic seeds’ hereafter) in the dietary spectra improves growth, we reared larvae of *S. saxatilis, L. equestris*, and *H. superbus* under four dietary treatments. Starting with pre-weighed first instar larvae, bugs were raised either on sunflower seeds only (positive control), a seed mixture comprising 11-15 natural host plant species to reflect the broad dietary spectra of *S. saxatilis* and *L. equestris* (Supplemental Table 1), the identical seed mixture supplemented with toxic seeds, or toxic seeds only. To establish species-specific seed mixtures, we selected natural host plant species based on literature data or own field observations. Toxic seeds were selected based on natural associations of the bug species with toxic plants in the field. Specifically, we used seeds of *D. purpurea* for *H. superbus*, seeds of *A. vernalis* for *L. equestris*, and seeds of *C. autumnale* for *S. saxatilis*.

Untreated, ripe seeds were either obtained commercially or collected in the field (*Digitalis purpurea*, Eberbach 2016). Seed mixtures consisted of 11 host species from 4 botanical families for *S. saxatilis* and 15 host species from 7 botanical families for *L. equestris*. Due to the lack of natural history data for *H. superbus*, we used the seed mixture used for *S. saxatilis* also for *H. superbus*. Within seed mixtures, we standardized proportions of individual plant species based on mass. Growth experiments were carried out in spatially randomized Petri dishes in a growth chamber (Binder KBWF 240, Tuttlingen, Germany) under the following conditions: 16: 8 h day/night cycle, 26 °C (*S. saxatilis*) or 28 °C (*L. equestris* and *H. superbus*), and 60 % humidity over the course of three weeks. We lined Petri dishes (60 mm x 15 mm, with vents, Greiner Bio-One, n = 11 for all species and diets) with filter paper and added three 1^st^ instar larvae to each dish. Petri dishes were supplied with a water source (see above) and 140.7 mg (± 0.72 SE, *S. saxatilis*), 145.1 mg (± 1.03 SE, *L. equestris*), or 140.1 mg (± 0.69 SE, *H. superbus*) seeds.

We recorded body mass weekly by anesthetizing all bugs of a Petri dish with CO2 and weighing them jointly. In addition, we recorded survival of bugs weekly. After the experiment, we transferred at least one bug from each Petri dish to fresh sunflower seeds for a period of two weeks to clean guts from potentially remaining toxins. Note that not all bugs had reached the adult stage at the time of transfer, thus the time spent on sunflower seeds during the adult stage varied and some bugs were still larvae at the time of analysis (see Supplemental Table 2 and the supplementary results). Finally, bugs were frozen at −80°C, freeze dried, weighed, and analyzed to quantify sequestered toxins via high performance liquid chromatography (HPLC) as described below (n = 11 for all species and diets, except of *H. superbus* on the seed mixture and pure *Digitalis* seeds and *L. equestris* on the seed mixture containing *Adonis* seeds, n = 10).

Growth of bugs on different seed mixtures, approximated as body mass (i.e. the total weight of all bugs per Petri dish divided by the number of remaining individuals) after three weeks of feeding, was analyzed by ANCOVA using the standard least squares method in JMP (JMP v. 13, SAS Institute, Cary, NC, USA). Samples sizes were 11 (i.e. 11 Petri dishes with up to three bugs) for all species and treatments except of *H. superbus* on the seed mixture and pure *D. purpurea* seeds (n = 10). Body masses were log_10_-transformed to achieve homogeneity of variances and normality of residuals. Dietary treatment was treated as a main effect, and initial mass of the bugs was included as a covariate. Two individuals of *L. equestris (Adonis* treatment) and of *H. superbus (Digitalis* and seed mixture plus *Digitalis* treatment) were statistical outliers, but their exclusion did not affect the direction or significance of effects. In addition to comparing final body mass after three weeks of feeding, we also modelled growth as a continuous process (see supplementary electronic material). We analyzed developmental time across treatments by comparing the number of days (log_10_-transformed) after which at least one bug per petri dish reached the adult stage using ANOVA in JMP (n = 11 for *L. equestris* on all treatments; n = 9 for *H. superbus* on the seed mixture without *Digitalis-seeds*, and n = 10 for all other treatments; note that some individuals turned into adults after being transferred to sunflower seeds). We omitted this analysis for *S. saxatilis* since on the *Colchicum* diet only one individual turned adult during the time of observation.

### Preparation of insect specimens for HPLC analysis

We determined the concentration of plant toxins sequestered by individual milkweed bugs with HPLC-DAD. We added 1 ml methanol (HPLC grade) containing 0.01 mg of the internal standard digitoxin (Sigma-Aldrich, Taufkirchen, Germany) to freeze-dried specimens of *L. equestris* and *S. pandurus*. For *H. superbus*, digitoxin was replaced by oleandrin (Phytolab, Vestenbergsgreuth, Germany) due to the natural occurrence of digitoxin in *Digitalis*. For specimens of *S. saxatilis* and *S. pandurus*, we used no internal standard and quantified colchicum alkaloids and bufadienolides with an external calibration curve (see below). After the addition of ca. 900 mg zirconia beads (Roth, Germany), specimens were homogenized in a Fast-Prep-24 homogenizer (MP Biomedicals, Germany) for two 45 sec cycles at a speed of 6.5 m/sec. After centrifugation at 16,100 x g for 3 min, supernatants were transferred to fresh 2 ml plastic vials (Sarstedt, Germany). Extractions were repeated once more with pure methanol. Pooled supernatants were evaporated under nitrogen gas. Subsequently, we re-suspended samples by adding 100 μl of methanol (200 μl for field-collected *S. pandurus* and *saxatilis*) and agitation in the Fast-Prep-24 homogenizer (45 s, 6.5 m/sec) without beads to facilitate dissolution of dried residues. Finally, samples were centrifuged (16,100 x g, 3 min) and filtered into HPLC-vials using Rotilabo®-syringe filters (nylon, 0.45 μm, Roth, Germany). Eggs obtained from *S. saxatilis* (n = 5; pools of 7, 18, 22, 4, and 7 eggs) females collected in Berghausen on May 5^th^ 2016 and from field-collected *L. equestris* (Lebus, n = 3; pools of 27, 28, and 63 eggs) were freeze dried and extracted as described above with 2 x 500 μl or 2 x 1 ml methanol, respectively. Details on harvesting of haemolymph and defensive secretion (i.e. clear droplets released at the integument upon attack) of *S. saxatilis*, as well as the preparation of dried museum specimens and plant seeds for chemical analysis are described in the electronic supplementary material.

### HPLC analysis of cardenolides, bufadienolides, and colchicum alkaloids

Fifteen microliters of extract were injected into an Agilent 1100 series HPLC and compounds were separated on an EC 150/4.6 NUCLEODUR^®^ C18 Gravity column (3 μm, 150 mm x 4.6 mm, Macherey-Nagel, Düren, Germany). Cardenolides and bufadienolides were eluted at a constant flow of 0.7 ml/min at 30°C with an acetonitrile–H2O gradient as follows: 0–2 min 16% acetonitrile, 25 min 70% acetonitrile, 30 min 95% acetonitrile, 35 min 95% acetonitrile, 37 min 16% acetonitrile, reconditioning for 10 min at 16% acetonitrile. We recorded UV absorbance spectra from 200 to 400 nm with a diode array detector. Peaks with symmetrical absorption maxima between 216 and 222 nm were interpreted as cardenolides, integrated at 218 nm and quantified based on the peak area of the known concentration of the internal standards digitoxin or oleandrin. Peaks with a symmetrical absorption maximum of 300 nm were interpreted as bufadienolides. For bufadienolide analysis, we used an external calibration curve based on proscillaridin A.

Colchicum alkaloids were eluted at a constant flow of 0.7 ml/min at 30°C with an acetonitrile–0.25 % phosphoric acid gradient as follows: 0–2 min 10% acetonitrile, 10 min 40% acetonitrile, 15 min 80% acetonitrile, 16 min 10% acetonitrile, reconditioning for 5 min at 10% acetonitrile. UV absorbance spectra were recorded from 190 to 400 nm by a diode array detector. Peaks with absorption maxima at 245 and 350 nm resembling the absorption spectra of colchicine were recorded as colchicosides and quantified at 350 nm. Colchicine-equivalents were calculated based on an external colchicine calibration curve.

Analysis of chromatograms was carried out with the Agilent ChemStation software (B.04.03). Details on the evaluation of individual datasets are described in the electronic supplementary material. Individual compounds were identified by comparisons of UV spectra and retention time with commercial reference compounds and liquid chromatography – mass spectrometry (details are described in the electronic supplementary material).

For analyzing sequestered toxins during seed feeding assays we used Welch’s test (JMP) and the Game-Howell post-hoc test (see www.biostathandbook.com) on log_10_ transformed data to evaluate differences across treatments, since data for *L. equestris* and *S. saxatilis* did not meet the assumption of equal variance. For this analysis, we excluded all data for individuals of *H. superbus* and *L. equestris* raised on sunflower seeds or seed mixtures without toxic seeds, as these lacked sequestered cardenolides.

### Behavioral assays with lacewings

We used larvae of the lacewing *Chrysoperla carnea* (Neuroptera, Chrysopidae) to test the effect of sequestered plant metabolites against an arthropod predator. Eggs of *L. equestris, H. superbus*, and *S. pandurus* were transferred to either sunflower seeds (non-toxic controls) or toxic seeds (*A. vernalis* for *L. equestris*, *D. purpurea* for *H. superbus*, and *U. maritima* for *S. pandurus*). Petri dishes were supplied with a water source as described above. After at least 3 (*S. pandurus*) to 5 (*H. superbus*) days in a growth chamber (Binder KBWF 240) at a 16: 8 h day/night cycle, 28 °C, 60% humidity, 1^st^ to 3^rd^ instar larvae (mainly 1^st^ and 2^nd^) were presented individually to a lacewing larva (2^nd^ or 3^rd^ instar). Larvae hatched from eggs of field-collected *S. saxatilis* were maintained on *C. autumnale* seeds for at least 4 days as described above. Due to the maternal transfer of colchicum alkaloids into the eggs, we used *O. fasciatus* larvae raised on sunflower seeds as a non-toxic negative control for these assays. We saved at least five individuals of each species from both treatments (toxic seeds and non-toxic sunflower seeds) to assess the amount of toxins sequestered using HPLC-DAD as described above.

Lacewing larvae were obtained commercially (Sautter & Stepper GmbH, Ammerbuch, Germany), maintained individually on *Sitotroga* (Lepidoptera: Gelechiidae) eggs (Katz Biotech AG, Baruth, Germany) for 3-5 days at room temperature and starved for two days before the experiments. In a first set of experiments, one milkweed bug larva was exposed to one lacewing larva in a Petri Dish (60 mm x 15 mm, with vents) and observed until the lacewing larvae attacked for the first time. After the first attack was over, lacewing larvae were removed and milkweed bug larvae were provided with a sunflower seed and water and checked for survival the next day. We excluded trials in which lacewing larvae did not attack bug larvae over the time of observation. The proportion of surviving milkweed bug larvae after the first attack by a lacewing across treatments was compared using two-tailed Fisher’s exact test in JMP.

In a second set of experiments, we assessed the effect of sequestered toxins on consumption of milkweed bug larvae by lacewing larvae. For *L. equestris*, both experiments (survival and feeding) were carried out twice since we initially assumed that the lack of an effect was due to the thick wall of the *Adonis vernalis* follicle rendering the seed inaccessible to small *L. equestris* larvae. Therefore, we repeated the experiments and maintained larvae on *A. vernalis* follicles chopped with a razor blade. Due to an overall lack of effect on lacewing behavior (see below), both experiments were combined for analyses. Details of this experiment are described in the electronic supplementary material.

### Behavioral assays with birds

Adults of *S. saxatilis* (specimens from Berghausen, see above), *L. equestris* and *H. superbus* (specimens from Eberbach, see above), either raised on sunflower seeds (control) or on toxic seeds (*C. autumnale* for *S. saxatilis*, *A. vernalis* for *L. equestris*, and *D. purpurea* for *H. superbus*), were offered to hand-reared juvenile great tits (*Parus major*; Passeriformes: Paridae) to test whether sequestered toxins protect bugs against avian predators. For this purpose, we raised first and second instar larvae to adults either on pure sunflower seeds or on a 1:1 mixture of sunflower with toxic seeds in plastic containers and supplied them with water as described above. Larvae were maintained in a growth chamber (Fitotron^®^ SGC 120, Weiss Technik, Loughborough, UK) at a 16: 8 h day/night cycle, at 26 - 27°C, and 60 % humidity. Before the experiment, we transferred bugs to pure sunflower seeds and maintained them at least for one week under the same conditions as described above to purge their guts from potentially retained plant toxins.

Birds were tested individually in wood-frame cages (70 × 70 × 70cm) with wire-mesh walls, equipped with a perch, a water bowl and a feeding tray. Cages were illuminated by daylight-simulating Osram Biolux 18-W/965 tubes. Prey was offered to birds in glass Petri dishes (diameter 50 mm) placed on the feeding tray. This way, all prey items appeared conspicuous against the light-colored background of the wooden tray. Before the experiment, the birds were habituated to the experimental cage, trained to eat mealworms from the tray, and deprived of food for 2 hours. We observed the birds through a one-way glass in the front wall of the cage and recorded their behavior using the Observer XT (Noldus) software. Details of hand-rearing juvenile birds are described in the electronic supplementary material.

Birds (100 in total) were divided into 3 experimental groups: 40 birds were tested with *S. saxatilis*, 40 birds with *L. equestris* and 20 birds with *H. superbus*. Within each group, half of the birds were tested with bugs raised on seeds of their respective toxic host plants and the other half with bugs raised on sunflower seeds (non-toxic control). To account for the effects of prey novelty on bird responses we used larvae of Jamaican field crickets (*Gryllus assimilis*) of a similar size as the bugs as a palatable prey control that would be unfamiliar to the birds. The experiment consisted of six five-minute trials (following immediately one after another) in which the birds were alternately offered a milkweed bug or a cricket, starting with the cricket. In each trial we recorded if the prey was attacked (pecked or seized), killed and eaten (at least partly), latency of the first attack, duration of prey handling, and number and duration of discomfort-indicating behavior (beak wiping and head shaking; supplementary results). If the bug was attacked but alive at the end of the trial, it was provided with water and sunflower seeds and checked for survival on the next day.

Bird predation data were analyzed using generalized linear models in R^53^. Attack rates and survival rates of the bugs were compared across the three trials using generalized estimating equation models (GEE, package geepack^54^) with binomial errors. Trial number, bug species and host plant toxicity were entered as fixed effects and bird individual (id) as a random effect. The models initially included all possible two-way interactions and were simplified by comparing nested models using the Quasi-information criterion (QIC). In the analysis of survival rates, only the data from bugs attacked in respective trials were included. To find out whether the general effect of host plant toxicity on reactions of birds also holds for each of the milkweed bug species studied, the abovementioned models were run separately for each bug species. Besides attack and survival rates, we also analyzed attack latencies, durations of discomfort-indicating behavior of birds, whether the bugs were (at least partly) eaten, and survival of bugs compared to control crickets; see the electronic supplementary material for details.

### Injection experiments to assess cardenolide and colchicine resistance

We injected adults of field-collected *Pyrrhocoris apterus*, or sunflower-raised *O. fasciatus, S. saxatilis* (Berghausen), and *S. pandurus* (Portugal) with colchicine or the cardenolide ouabain to test for the ability to tolerate these toxins in the body cavity (i.e. to mimic sequestration). We injected one microliter of toxins (dissolved in phosphate buffered saline, PBS, pH 7.4) or PBS as a control with glass capillary needles. Solutions were injected laterally between the penultimate and the last abdominal segment using a micromanipulator and a microsyringe pump injector (World Precision Instruments, Sarasota, FL, USA) under a dissecting microscope (n = 8 individuals per dose).

In total, we carried out three injection experiments. In a first experiment, *P. apterus*, *O. fasciatus*, and *S. saxatilis* were injected with either a high dose of ouabain (5 mg/ml) or colchicine (10 mg/ml, Figure 4). As we observed no effect of colchicine in *S. saxatilis* as opposed to the other species, we injected *S. saxatilis* with an even higher dose of colchicine in a second experiment and again injected 5 mg/ml ouabain in additional specimens for comparison. Since *P. apterus* responded to both toxins in the first trial, we injected 0.1, 1, 5, or 10 mg/ml colchicine or 0.1, 1, 2.5, 5 mg/ml ouabain (i.e. 0.1 to 10 μg/individual and 0.1 to 5 μg/individual, respectively) within the same attempt to address the extent of resistance quantitatively. *O. fasciatus* that tolerated 5 mg/ml ouabain were only injected with increasing concentrations of colchicine (0.1, 1, 5 and 10 mg/ml, i.e. 0.1 to 10 μg/individual). To keep the number of injected animals as low as possible we omitted injections of PBS during experiment 2 (tolerance to injections per se was already apparent from the first experiment). Lastly (experiment 3), we injected *S. pandurus*, a congener of *S. saxatilis* with PBS, 0.1, 1, 5, or 10 mg/ml colchicine. After injection, we maintained bugs individually in Petri dishes with a water source (see above) and one sunflower seed under ambient conditions. On the next day, we assessed bugs for signs of paralysis (i.e. inability to walk, slowed movement of legs and antennae). Injection assays with one dose of a toxin were analyzed using Fisher’s exact test in JMP with Bonferroni correction for multiple comparisons. For comparing dose-dependent effects, we used the Cochrane-Armitage trend test in JMP.

### Statistical analyses

For better readability, statistical analyses are explained at the end of each methods section (see above). If not mentioned in the main text, sample sizes are reported in the figure legends.

## Results

### Growth of milkweed bugs on different diets

We tested if the availability of ‘toxic seeds’ influences larval development. After three weeks, a diet of pure sunflower seed resulted in maximal growth in all three insect species tested (Figure 2a-c). We found a significant effect of seed diet on growth of *L. equestris* (F_3,39_ = 56.431, p < 0.001) and *S. saxatilis* (F_3,39_ = 56.631, p < 0.001), but not of *H. superbus* (F_3,37_ = 0.425, p = 0.736) which grew equally across all treatments. On diverse seed mixtures without toxic seeds, *S. saxatilis* grew as well as on sunflower seeds (LSMeans Tukey HSD: p = 0.883) while *L. equestris* only reached about half of the body mass compared to the sunflower diet (LSMeans Tukey HSD: p < 0.001). Inclusion of ‘toxic seeds’ into these mixtures did not increase growth for either of the species (LSMeans Tukey HSD: *L. equestris*, p = 0.384; *S. saxatilis*, p = 0.936). In fact, diets consisting exclusively of toxic seeds resulted in lower body mass compared to seed mixtures for both species (LSMeans Tukey HSD: *L. equestris*, p < 0.001; *S. saxatilis*, p < 0.001), and in a reduction of > 50 % in body mass compared to the sunflower diet (LSMeans Tukey HSD: *L. equestris*, p < 0.001; *S. saxatilis*, p < 0.001). Initial body mass affected the final weight of *S. saxatilis* (F_1,39_ = 5.453, p = 0.025) but not of *L. equestris* (F_1,39_ = 0.034, p = 0.855) and *H. superbus* (F_1,37_ = 0.051, p = 0.823). The modelling of larval growth as a continuous process and comparison of absolute growth rates revealed similar results (Supplemental Figure 1. We found negative effects of *Adonis* seeds on developmental time in *L. equestris* but not of *Digitalis* seeds in *H. superbus* (supplementary results).

**Figure 2.**
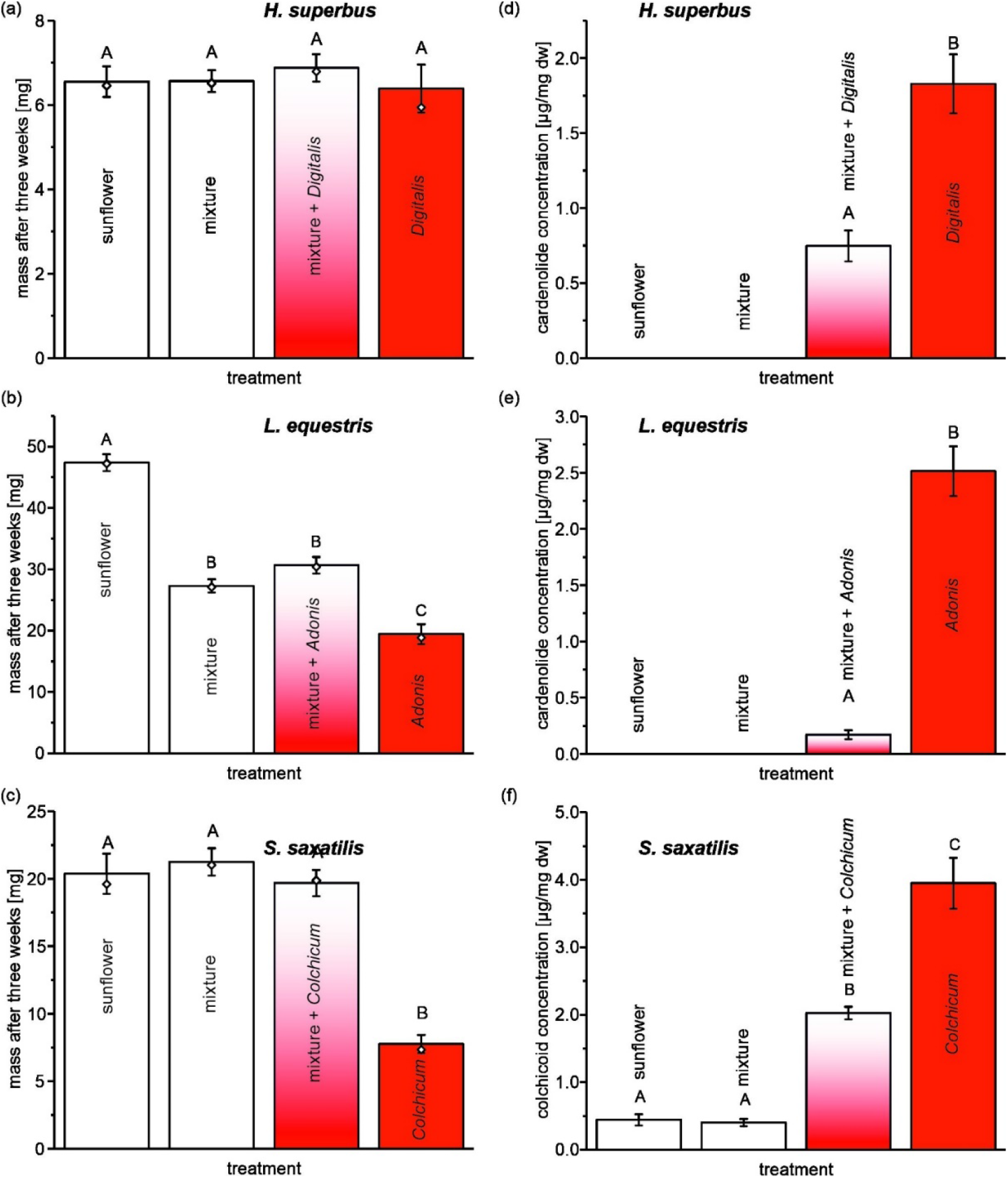
Growth of milkweed bugs and sequestration of plant toxins across different diets. Larvae of *H. superbus* (a), *L. equestris* (b), and *S. saxatilis* (c) were raised on sunflower seeds, a seed mixture, the same mixture containing either *Digitalis*- (*H. superbus*), *Adonis*- (*L. equestris*), or *Colchicum*- (*S. saxatilis*) seeds, or pure *Digitalis*-, *Adonis*-, or *Colchicum-see*. Larval mass was recorded over a period of three weeks. Bars represent means of body mass after three weeks ± SE. Diamonds represent retransformed model means of data that were log_10_ transformed for statistical analysis. Sample sizes for the obtained body masses for *H. superbus* were: sunflower = 11, mixture = 10, mixture plus *Digitalis* = 11, *Digitalis* = 10; for *L. equestris* and *S. saxatilis*: n = 11 for all diets. After the growth experiment, bugs were harvested for chemical analyses. The amount of sequestered cardiac glycosides from *Digitalis* (d) and *Adonis* (e) or colchicum alkaloids (f) was always highest on pure diets but sequestration was also substantial in seed mixtures containing *Digitalis*-, *Adonis*-, or *Colchicum*-seeds. Colchicum alkaloids found in bugs raised on sunflower seeds or seed mixtures lacking *Colchicum*-seeds originate from maternal egg transfer. Bars represent mean concentrations of sequestered toxins ± SE. Sample sizes are identical to the ones mentioned above except of n = 10 for *L. equestris* on the seed mixture with *Adonis*. Different letters above bars indicate significant differences among treatments (p < 0.05).

### Sequestration in milkweed bugs on different diets

In addition to growth, we also quantified sequestration of cardenolides and colchicum alkaloids. We found substantial sequestration of cardenolides in *H. superbus* and in *L. equestris* raised on pure *Digitalis* or pure *Adonis* seeds and seed mixtures containing toxic seeds, respectively, while bugs raised on sunflower seeds or seed mixtures without toxic seeds were devoid of cardenolides (Figure 2d,e). Similarly, *S. saxatilis* raised on seeds of *Colchicum* and seed mixtures containing *Colchicum* seeds sequestered high amounts of colchicum alkaloids (Figure 2f). Bugs from diet treatments lacking *Colchicum* seeds also contained low levels of these toxins which are most likely derived from field-collected females by transfer via the egg. The amount of sequestered toxins was always highest on the diets comprised of toxic seeds only (Welch’s test of diet effect; *H. superbus:* F_1,19_ = 18.863, p < 0.001; *L. equestris:* F_1,10_ = 66.206, p < 0.001; *S. saxatilis:* F_3,20_ = 83.568, p < 0.001).

### Sequestration of plant toxins in field-collected milkweed bugs

We collected adult milkweed bugs in the field to test for sequestration of plant toxins under natural conditions (Supplemental Figure 6). All specimens of *H. superbus* (n = 10) from a habitat with *D. purpurea* contained cardenolides (Figure 1c) ranging from 1 to 8.9 μg per mg dry mass (2.6 to 28.6 μg per insect, n = 10). *H. superbus* collected from a *Digitalis-free* habitat (n = 12) that we observed feeding on pods of *E. crepidifolium* invariably contained high amounts of sequestered cardenolides (Figure 1c) ranging from 23 – 61.2 μg/mg (41 – 147 μg per specimen). All *L. equestris* obtained from the *A. vernalis* site had cardenolide concentrations ranging from 0.14 to 28.12 μg/mg dry mass (4.7 to 459.6 μg per individual, n = 12; Figure 1b). Moreover, we detected cardenolides in eggs laid by field-collected females (0.23 μg/mg dry weight ± 0.01, n = 3; mean ± SE). *S. pandurus* collected from infructescences of *U. maritima* contained up to 17.8 μg bufadienolides per mg dry mass (up to 808 μg per individual, n = 7; Figure 1b).

Adults of *S. saxatilis* obtained from two different populations consistently contained colchicum alkaloids (Figure 1a) ranging from 0.05 – 6.2 μg/mg dry mass (1.55 – 113.7 μg per individual, Nüstenbach, n = 16) and 6.4 – 9.4 μg/mg dry mass (134.3 – 214.3 μg per individual, Berghausen, n = 9). Eggs from field-collected females contained colchicum alkaloids (0.94 μg/mg dry weight ± 0.26, n = 5; mean ± SE). Concentrations of colchicum alkaloids in *S. saxatilis* defensive secretion were more than 50 times higher compared to haemolymph (paired t-test on six paired samples: t = 4.234, df = 5, p = 0.008, Supplemental Figure 4). The comparison of sequestered toxins to toxins present in the host plant seeds revealed a clear overlap of individual compounds only in some insect species (Supplemental Figure 3) indicating extensive metabolism or selective uptake and up-concentration by the other species (see the electronic supplementary material for details on structural identification and comparison to seed extracts).

### Colchicum alkaloids in museum specimens

We screened 30 museum specimens of *S. saxatilis* from 21 locations in 10 European countries and one location in North Africa (Supplemental Figure 5). Although some of the specimens were more than 110 years old, we detected substantial amounts of colchicum alkaloids ranging from 0.8 μg to 182.5 μg per individual (58 ± 8.69, mean ± SE) in all specimens. Remarkably, only two of the specimens tested contained trace amounts of putative cardenolides, suggesting that sequestration of cardenolides does not play a role for the defense of *S. saxatilis*.

### Effect of sequestered toxins on lacewing predation

We assessed the effect of sequestered plant toxins on lacewing predation in four species of milkweed bugs. *H. superbus* larvae raised on seeds of *Digitalis* contained 3.15 ± 0.99 μg cardenolides per mg dry weight (mean ± SE, n = 7) and survived lacewing attacks more often compared to sunflower-raised individuals (Figure 3a; p = 0.02, two tailed Fisher’s exact test). However, even though *L. equestris* larvae raised on *Adonis* seeds contained higher cardenolide amounts (whole seeds: 4.95 ± 0.27 μg/mg, n = 8; chopped seeds: 5.01 ± 0.45 μg/mg, n = 5; mean ± SE), only one sunflower-raised larva out of a total of 63 *L. equestris* larvae tested (n = 32 for sunflower, n = 31 for *Adonis* seeds) survived the first attack by a lacewing larva (Figure 3b; p = 1, two tailed Fisher’s exact test). Similarly, *S. pandurus* larvae raised on *Urginea* seeds contained 16.11 ± 4.39 μg/mg bufadienolides (n = 7), yet only one *Urginea*-raised larva out of all 40 tested survived the first lacewing attack (n = 20 for both treatments, p = 1, two tailed Fisher’s exact test). In contrast, sequestration of colchicum alkaloids protected *S. saxatilis* from lacewing attacks. Thirteen out of 20 larvae derived from eggs of field-collected females survived the first attack by a lacewing larva (Figure 3c) while there was no survival in sunflower raised *Oncopeltus* larvae used as a control (p < 0.001, two tailed Fisher’s exact test). Concentrations of colchicum alkaloids in *S. saxatilis* larvae were 53.15 ± 3.12 μg/mg (n = 10, mean ± SE). Even though only two out of four systems tested showed an effect on survival, there was at least some evidence for a negative effect of sequestered compounds on consumption and weight gain by lacewing larvae in all four systems (Supplemental Figure 7, see the electronic supplementary material for details).

**Figure 3.**
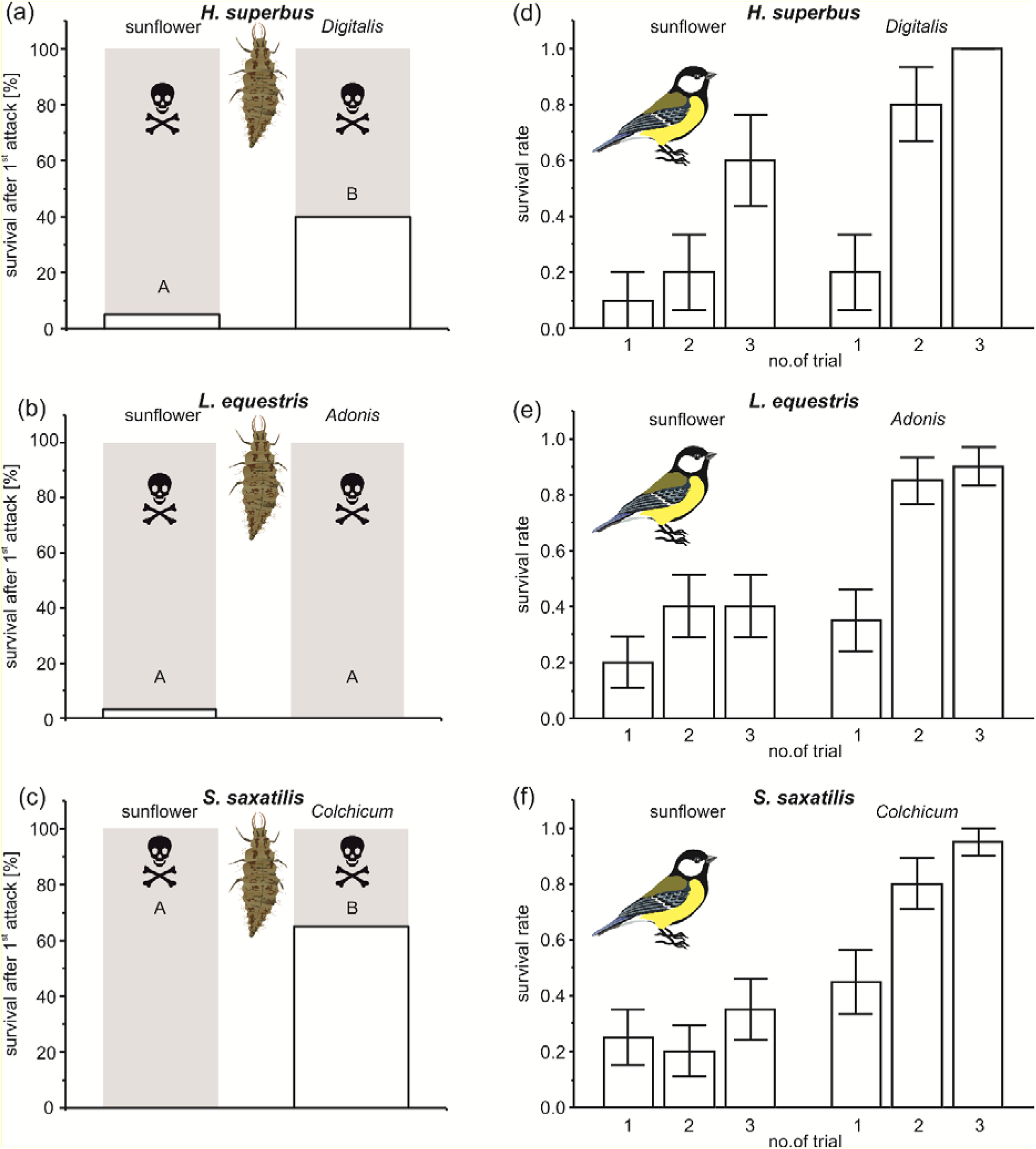
Predation of lacewing larvae (*C. carnea*) and great tits (*P. major*) on three species of milkweed bugs. Milkweed bugs were either raised on sunflower seeds (controls) or on seeds of the following plant species: *D. purpurea* (*H. superbus*), *A. vernalis* (*L. equestris*), or *C. autumnale* (*S. saxatilis*). Please note that in the experiment with *S. saxatilis* and lacewing larvae, sunflower raised larvae of *O. fasciatus* were used as control. Left panel: we assessed the proportion of milkweed bug larvae that survived the first lacewing attack. (a-c): survival of *H. superbus* (n = 20 for both diets), *L. equestris* (sunflower: n = 32, *A. vernalis*: n = 31), and *S. saxatilis* (sunflower: n = 20, *C. autumnale:* n = 20). Grey (skull) is the proportion of bugs, which were killed, white is the proportion of bugs, which survived the first attack by the lacewing larva. Different letters above bars indicate significant differences among treatments. Right panel: Survival rates of adult milkweed bugs across three successive encounters with juvenile *P. major*. (d-f): survival of *H. superbus, L. equestris*, and *S. saxatilis*. Only the data from bugs attacked by birds in respective trials are included. Bars represent mean survival rates ± SE.

### Effects of sequestration on defence against avian predators

We analysed the effects of sequestration of host plant toxins on defence against avian predators in *L. equestris, S. saxatilis* and *H. superbus*. Overall attack rates were similar for all bug species (GEE, χ^2^_2_ = 3.242, *P* = 0.198), but significantly lower when the birds were tested with bugs from toxic host plants (GEE, χ^2^_1_ = 5.596, *P* = 0.018). Attack rates decreased over the three successive trials (GEE, χ^2^_1_ = 115.785, *P* < 0.001), and the decrease was affected neither by bug species (GEE, trial: species interaction, χ^2^_2_ = 2.032, *P* = 0.362) nor host plant toxicity (GEE, trial: host plant interaction, χ^2^_1_ = 0.815, *P* = 0.367). However, a difference in this decrease of attack rates became apparent if species were analysed separately. In *S. saxatilis*, attack rates decreased over trials (GEE, χ^2^_1_ = 55.968, *P* < 0.001), and more so when the bugs were coming from *Colchicum autumnale* than from sunflower (GEE, hostplant, χ^2^_1_ = 2.414, *P* = 0.121; trial: host plant interaction, χ^2^_1_ = 6.009, *P* = 0.014). Likewise, attack rates towards *H. superbus* decreased over trials (GEE, χ^2^_1_ = 39.327, *P* < 0.001), and the decrease was steeper for bugs raised on *Digitalis purpurea* (GEE, host plant, χ^2^_1_ = 1.250, *P* = 0.412; trial: host plant interaction, χ^2^_1_ = 6.525, *P* = 0.011). Contrastingly, attack rates towards *L. equestris* decreased over trials irrespectively to host plant toxicity (GEE, trial, χ^2^_1_ = 28.957, *P* < 0.001; hostplant, χ^2^_1_ = 0.056, *P* = 0.813; trial: host plant interaction, χ^2^_1_ = 1.067, *P* = 0.302).

Overall survival rates following attack were higher in bugs from toxic host plants (GEE, χ^2^_1_ = 24.477, *P* < 0.001) and increased significantly over successive trials (GEE, χ^2^_1_ = 13.581, *P* < 0.001), with this increase being steeper in the bugs from toxic host plants (GEE, trial: host plant interaction, χ^2^_1_ = 9.341, *P* = 0.002; Figure 4d-f). Bug species affected neither the overall survival (GEE, χ^2^_2_ = 0.777, *P* = 0.678) nor its increase over the trials (GEE, trial: species interaction, χ^2^_1_ = 3.403, *P* = 0.183). Separate analyses for each species revealed that survival rate was generally higher if milkweed bugs were raised on toxic host plants (GEE, *S. saxatilis:* χ^2^_1_ = 12.471, *P* < 0.001; *L. equestris:* χ^2^_1_ = 10.037, *P* = 0.002; *H. superbus:* χ^2^_1_ = 5.086, *P* = 0.024), and it increased over the trials (GEE, *S. saxatilis:* χ^2^_1_ = 10.911, *P* < 0.001; *L. equestris:* χ^2^_1_ = 5.644, *P* = 0.018; *H. superbus:* χ^2^_1_ = 17.788, *P* < 0.001; Figure 4d-f). The increase was steeper in *S. saxatilis* and *L. equestris* coming from toxic host plants (GEE, trial: host plant interaction, *S. saxatilis:* χ^2^_1_ = 6.097, *P* = 0.014; *L. equestris:* χ^2^_1_ = 4.394, *P* = 0.036), but there was no significant difference in *H. superbus* (GEE, χ^2^_1_ = 2.442, *P* = 0.118).

**Figure 4.**
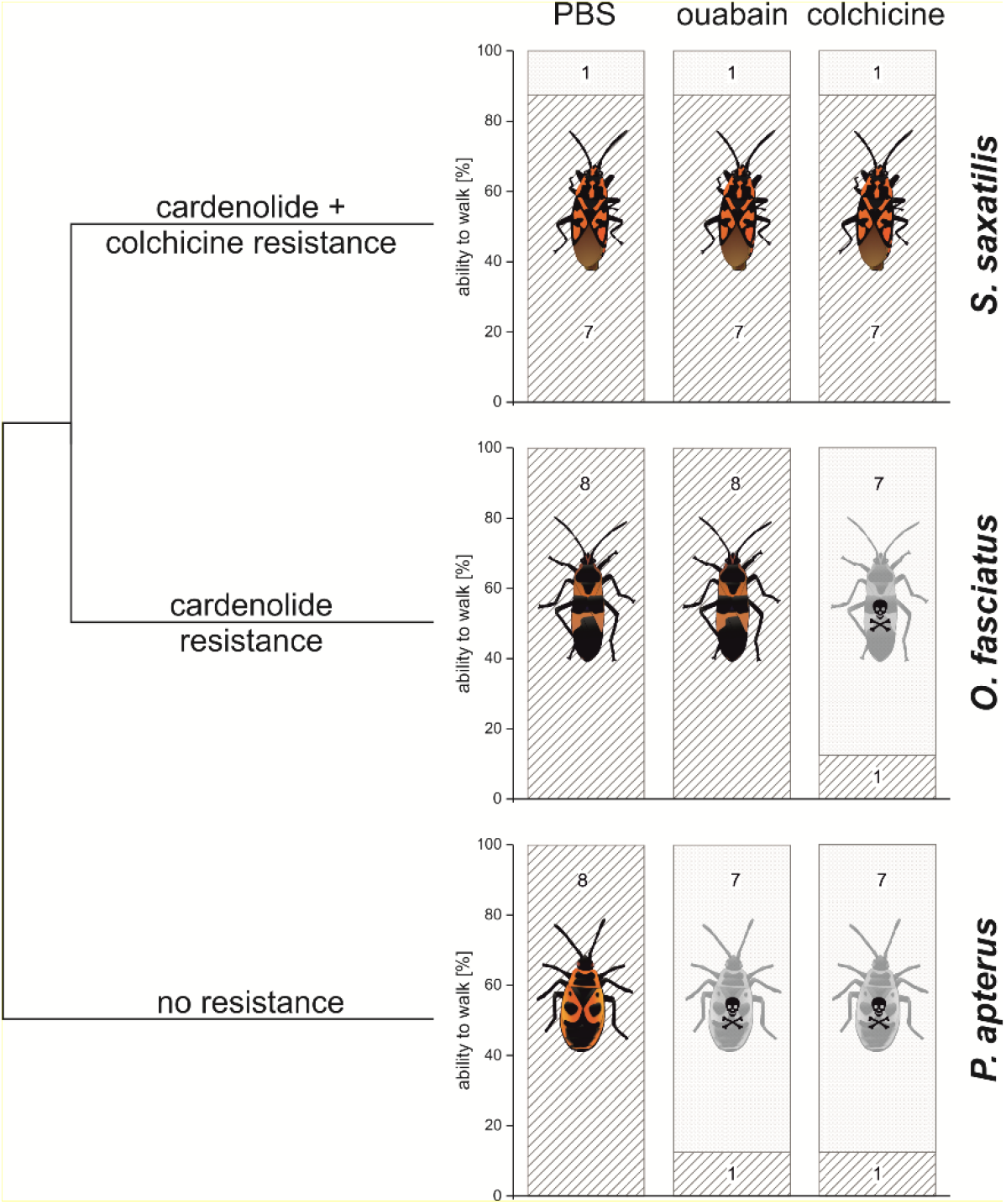
Resistance of *P. apterus, O. fasciatus*, and *S. saxatilis* to injected toxins. Adult hemipteran specimens were injected with either PBS (control), the cardenolide ouabain (5 mg/ml, i.e. 5 μg/individual), or the alkaloid colchicine (10 mg/ml, i.e. 10 μg/individual). On the left, we mapped resistance phenotypes on a scheme representing the phylogenetic relationships of the species involved. Bar charts on the right show the percentage of individuals that showed no signs of intoxication at the next day after injecting toxins (hatched). Insect icons are intended to visualize either a toxic (grey bug with skull) or no toxic effect (colored bug). Numbers in stacked bars represent the actual number of affected or unaffected individuals. Note that we also tested *S. pandurus*, a congener of *S. saxatilis*, and found it not to possess resistance to colchicine (see Supplemental Figure 10).

In addition, sequestered plant toxins increased bird attack latencies across trials (Supplemental Figure 8), the duration of discomfort-indicating behavior (Supplemental Figure 9), and decreased the chance that the birds would consume the bugs. Nevertheless, compared to crickets, milkweed bugs devoid of sequestered toxins were not entirely undefended against the birds, emphasizing the role of endogenous scent-gland secretion (see the electronic supplementary material for details).

### In vivo tolerance to injected ouabain and colchicine

Due to the rare occurrence of colchicine in nature which is restricted to *Colchicum* spp. and other Colchicaceae^55^, we predicted that *S. saxatilis* was not preadapted to this toxin but evolved novel resistance traits against colchicine. In addition, we predicted this species to retain resistance against cardiac glycosides based on its evolutionary history. To test for toxin resistance and to mimic sequestration, we injected colchicine or ouabain directly into the body cavity of milkweed bugs. Besides *S. saxatilis*, we used *Pyrrhocoris apterus* (Pyrrhocoridae) as a non-adapted outgroup, and the lygaeine *Oncopeltus fasciatus* as a cardenolide-sequestering milkweed specialist. None of the species tested was affected by blank injections of the solvent PBS (Figure 4, Supplemental Figure 10). As expected, *P. apterus* was unable to tolerate injections of either 5 μg ouabain or of 10 μg colchicine (ouabain vs. PBS, p = 0.001; colchicine vs. PBS, p = 0.001; two tailed Fisher’s exact test at p = 0.025 after Bonferroni correction). All *O. fasciatus* individuals tolerated an injection of 5 μg ouabain but were not able to tolerate 10 μg colchicine (p = 0.001, two tailed Fisher’s exact test). As predicted, *S. saxatilis* tolerated injections with both classes of toxins (ouabain vs. PBS, p = 1; colchicine vs. PBS, p = 1; two tailed Fisher’s exact test at p = 0.025 after Bonferroni correction). We furthermore found that *S. pandurus*, a congener of *S. saxatilis*, was not able to tolerate colchicine injections. Moreover, *O. fasciatus* responded to colchicine and *P. apterus* to ouabain and colchicine, in a dose-dependent manner (Supplemental Figure 10). *S. saxatilis* tolerated injections of up to 30 μg colchicine per animal, the highest dose tested (see the electronic supplementary materials for details).

## Discussion

It is widely accepted that coevolution between insects and plants occurs in a multitrophic context. Nevertheless, our understanding of the evolutionary drivers is still limited especially with regard to the underlying mechanisms. Coevolutionary theory posits that occupation of novel dietary niches depends on insect resistance to host plant toxins and that resistance traits may interfere with dietary breadth. Recently, we have shown that in addition, interactions with higher trophic levels (predators and parasitoids) are likely to select for specific resistance traits in insects^22^. However, the interplay of specific adaptations and acquisition of plant toxins for defense as drivers of host shifts^5^ and specialization has never been addressed. Here, we tested if sequestration of plant toxins and preadaptation in milkweed bugs mediated specific associations with particular host plants. Detailed analyses of the selective forces directing the evolution of insect-plant interactions are mandatory to unravel the function of ecosystems not only for basic research but also for conservation efforts and the advancement of coevolutionary theory.

In accordance with their global association with plants in the Apocynaceae, milkweed bugs (Lygaeinae) are preadapted to sequester cardiac glycosides. We identified three milkweed bug species, *L. equestris*, *H. superbus*, and *S. pandurus*, which independently colonized plants from four botanical families (Asparagaceae, Brassicaceae, Plantaginaceae, and Ranunculaceae) that produce cardiac glycosides convergently, and found that all three species sequester these toxins from their evolutionarily novel hosts. A fourth species, *S. saxatilis*, has furthermore evolved the ability to sequester alkaloids from *C. autumnale* (Colchicaceae), likely using some of the same mechanisms for uptake, storage and release as are used for cardiac glycoside sequestration.

Across the three systems tested (*H. superbus* on *D. purpurea, L. equestris* on *A. vernalis*, and *S. saxatilis* on *C. autumnale*) we did not find improved growth when toxic seeds of the respective novel hosts were included into seed mixtures. On pure diets of toxic *A. vernalis* and *C. autumnale* seeds, growth was reduced substantially, indicating that seeds from these plant species alone are not a suitable diet. Only *H. superbus* grew equally well on toxic *D. purpurea* seeds as on all other diets, suggesting a higher degree of dietary specialization for this species. Our results demonstrate that seeds from toxic host plants are not required for successful development, which is supported by the fact that species such as *H. superbus* and *L. equestris* are easy to maintain in the laboratory exclusively on sunflower seeds over many generations without an apparent effect on fitness. While toxic host plants may also provide important nutritional resources temporarily, it seems unlikely that specific associations with these plants were evolutionarily driven by the benefit of occupying novel dietary niches. This is in line with several milkweed bug species being generalist seed predators that can feed on a tremendous variety of host plant species.

Throughout its distributional range, *L. equestris* is closely associated with *Vincetoxicum hirundinaria*, an Apocynaceae that is lacking cardiac glycosides but provides *L. equestris* with an unknown defense^29^. Nonetheless, *L. equestris* reached smaller body size when raised on *V. hirundinaria* compared to being raised on sunflower seeds^29^ and other fitness related traits such as fertility, mortality, and developmental time were not different across the two diets^56^. In nature, both *L. equestris* and *S. saxatilis* in fact use a great diversity of host plants. For *L. equestris*, 60 plant species from roughly 20 botanical families^45^ and for *S. saxatilis* more than 40 species from over 15 families have been recorded (see Supplemental Table 3). Consequently, both species can be considered generalists from a dietary perspective.

We have shown earlier that cardiac glycoside resistant Na^+^/K^+^-ATPases and sequestration of cardiac glycosides apparently are synapomorphic traits of the Lygaeinae^40^. In addition, milkweed bugs concentrate cardiac glycosides far above haemolymph levels in specialized storage compartments^33,34^ from where they are released upon predator attack. Remarkably, we found an identical suite of adaptations to colchicum alkaloids in *S. saxatilis* with colchicine and related alkaloids being highly enriched in the defensive secretion compared to the haemolymph. To tolerate sequestration of these compounds, *S. saxatilis* evolved a novel resistance trait against colchicine that is not present in its congener *S. pandurus*.

Specialization of *S. saxatilis* to *C. autumnale* (and maybe other *Colchicum* species) is evidenced by our screening of museum specimens. The presence of colchicum alkaloids in 30 randomly selected specimens from 11 countries in Europe and North Africa clearly shows that each individual accessed *Colchicum* during its lifetime, while the lack of cardenolides suggests that *S. saxatilis* completely shifted from the use of cardenolides to the novel defense. Oviposition into *Colchicum* seedpods and allocation of high amounts of alkaloids into the eggs finally supports a close association of *S. saxatilis* with *C. autumnale*. Remarkably, *S. saxatilis* still maintains resistance to cardiac glycosides and accumulated resistance traits against different classes of plant toxins over evolutionary time, even though target site insensitivity of Na^+^/K^+^-ATPase was suggested to incur a physiological cost in *O. fasciatus*^42^.

The results of our predation assays revealed that feeding on either cardiac glycoside- or colchicum alkaloid-containing seeds at least partially protects milkweed bugs against lacewing larvae and passerine birds. We found higher survival after lacewing attacks for *H. superbus* raised on *D. purpurea* seeds, and for *S. saxatilis* larvae that derived colchicum alkaloids from their eggs. In contrast, *L. equestris* raised on *A. vernalis* seeds and *S. pandurus* raised on *U. maritima* seeds were not protected against lacewings, suggesting that predator-prey interactions are likely affected by the source and quality (rather than quantity) of sequestered plant toxins, the sequestering insect species, or a combination of both. Nevertheless, together with previous studies^32,57^ our results suggest that sequestration of plant toxins mediates effective defense of milkweed bugs against arthropod predators.

In experiments with avian predators, sequestration of host plant chemicals decreased bird attack rates and increased prey survival compared to sunflower-raised bugs. This effect was present in all three milkweed-bug species, colchicine being equally effective as cardenolides. Aversiveness of sequestered chemicals was also evidenced by discomfort-indicating behavior following contact with bugs from toxic host plants. Higher effectiveness of sequestered than autogenous chemicals against avian predators has also been found in other studied systems, e.g. leaf beetles^58^ and lanternflies^59^. Nevertheless, our results show that the autogenous scent-gland secretion alone still increases bug survival compared to undefended prey. Our findings indicate that besides being highly toxic, cardenolides^60^ and colchicine may protect prey due to an aversive taste. Consequently, the bugs derived from toxic host plants frequently survived bird attacks and were almost never eaten, while sunflower-raised bugs were frequently killed, and at least partly consumed.

The increased effectiveness of sequestration over autogenous secretion indicates that milkweed bugs represent an instance of automimicry^61^, i.e. interspecific variation in antipredatory defence when less defended or undefended individuals gain protection by resembling their better defended conspecifics. Decreasing attack rates and increasing bug survival across trials suggest that birds combined decisions based on visual cues with taste-sampling^62,63^, which allows to discriminate between defended individuals and automimics^64^ and taste-reject only the prey that actually contains toxins^65^. The reason why milkweed bugs maintain variation in host plant utilization and chemical defense could be a trade-off between development and defense^66^ mediated by toxic host plants representing a suboptimal food source. Nevertheless, different defense chemicals could still be equally effective against some predators^29^. Unpredictability of defense is also considered aversive by itself^67^, and can increase effectiveness of avoidance learning^68^. A broad spectrum of host plants in many milkweed-bug species suggests that in this taxonomic group automimicry is evolutionary stable^69,70^. Intraspecific variation in antipredatory defence of milkweed bugs may also affect their mimetic relationships with similarly colored prey species.

Sequestration of cardiac glycosides from *A. vernalis, D. purpurea, E. crepidifolium*, and *U. maritima* was most likely facilitated by preadaptation, yet sequestration of colchicum alkaloids is an entirely novel ability. However, while colchicum alkaloids drastically differ from cardiac glycosides in their mode of action or target site, the two types of compounds nonetheless share some similarities: both comprise small, chemically stable, and mostly lipophilic molecules that do not require enzymatic activation to become highly toxic. Therefore, even though *S. saxatilis* likely lacked preadaptations for tolerance of the novel toxin, its aposematism and sequestration machinery for uptake, specialized storage and release of toxins^33^ may well have facilitated the evolution of colchicum alkaloid sequestration. Our findings therefore demonstrate that host shifts of specialized insects can be mediated by preadaptation to specific toxins and convergent evolution of plant toxins in unrelated plant taxa. At the same time, we propose that suites of traits involved in sequestration of one type of chemical may similarly represent preadaptations facilitating shifts to entirely novel classes of chemically unrelated compounds, particularly if favored by large benefits (i.e. sequestration of highly toxic colchicine).

In conclusion, specialization in milkweed bugs is not an evolutionary dead end^4^, and evolutionary plasticity is maintained by different mechanisms including preadaptation with regard to different traits of the same syndrome. Our findings demonstrate that it is insufficient to classify the degree of specialization in insects solely based on their trophic interactions. Species that classify as dietary generalists may still specialize to host plants serving as a source of sequestered toxins. Interactions driven by the third trophic level (predators and parasitoids) can therefore direct specialization of bitrophic interactions between herbivores and their hostplants.

## Acknowledgments

We thank Michael Falkenberg for drawing our attention to *Spilostethus saxatilis* emerging from seedpods of *Colchicum autumnale*, Susanne Dobler for providing a lab strain of *Oncopeltus fasciatus*, Andreas Berger who observed *H. superbus* feeding on *E. crepidifolium* and shared the location with us, and Luis Vivas for supporting our search of *S. pandurus* feeding on *U. maritima*. The Museum für Naturkunde Berlin, the Senckenberg Deutsches Entomologisches Institut Müncheberg, and the Staatliches Museum für Naturkunde Karlsruhe, Germany, provided dry specimens of *S. saxatilis* for chemical extraction. We furthermore thank all the collectors of specimens and Hermann Falkenhahn for identifying host plants of *S. saxatilis*. We thank Anurag Agrawal for comments on our manuscript. Moreover, we thank the Junta Andalucia, Spain, the Landesamt für Umwelt Brandenburg, the Regierungspräsidium Karlsruhe, and the Struktur- und Genehmigungsdirektion Nord, Koblenz, Germany for issuing collecting permits for bugs and plant material. This work was supported by DFG grant PE 2059/3-1 to GP and the LOEWE program of the State of Hesse to AV and GP via funding the LOEWE Center for Insect Biotechnology & Bioresources and a Czech Science Foundation Grant 19-09323S to AE.

## Supplementary Materials

### Supplementary Methods

#### Modelling of milkweed bug growth in seed mixture experiments

In addition to comparing final body mass of milkweed bug larvae after three weeks of feeding, we also modelled larval growth as a continuous process using the four sequential body mass measurements recorded during the experiment (initial mass, mass at weeks 1, 2, and 3). Log-transformed body masses were modelled using an asymptotic regression model^71,72^, implemented as the *SSasymp* function in the *nlme* package for the statistical software R. Body mass was thus modelled as a function of the initial mass, a rate of increase, and an asymptotic mass at the end of the experiment. Effects of seed mixture were included for the rate of increase and asymptote parameters, while individual bugs (i.e. body mass means per petri dish) were treated as random effects to account for repeated measures. To compare growth between seed mixture treatments, the absolute growth rate of bugs (AGR, mass gain per day) was calculated for the final day of the experiment^72^.

#### Maintenance of milkweed bugs on sunflower seeds to purge toxins from guts

After the mixed seed feeding experiments, bugs were transferred to fresh sunflower seeds for 14 days to clean guts from residual dietary toxins that would bias the estimate of compounds sequestered into the body tissues. Since several true bug species, including *Oncopeltus fasciatus* are known to have a discontinuous digestive tract until the adult stage^73^, we cannot rule out that larvae had toxins remaining in their guts at the time of analysis (see Supplemental Table 2 for the number of days individual specimens had for purging during the adult stage). As an estimate, we quantified colchicum alkaloids in the gut of last instar larvae of *S. saxatilis* (extracted in 2 x 1 ml methanol, resuspended in 100 μl methanol and analyzed as described above) that were raised on *C. autumnale* seeds from the second larval instar and found them to contain only 1.44 μg alkaloids per gut (n = 4, SE = 0.76). Similarly, other authors found that the amount of cardenolides in filled guts of adult *O. fasciatus* raised on milkweed seeds was below a detection limit of 10 μg^74^. Consequently, it seems unlikely that substantial amounts of non-sequestered toxins remaining in the larval gut would have biased our results.

#### Comparison of colchicum alkaloids in haemolymph and defensive secretion of S. saxatilis

To compare concentrations of sequestered colchicum alkaloids between haemolymph and defensive secretion (i.e. clear droplets released at the integument upon attack), we used adults of *S. saxatilis* collected in the field (Berghausen, August 2016). Before collecting samples, we maintained bugs on pure *Colchicum* seeds (supplied with water) for 13 days under ambient conditions. To obtain defensive secretion, we anaesthetized bugs with CO2 and squeezed them between the blades of broad tweezers. We collected emerging droplets of clear defensive fluid using 0.5 μl glass capillaries. Next, we cut off a hind-leg at the femur to collect haemolymph. We determined the collected volume based on the filling level of the capillaries. In total, we obtained 13 samples of haemolymph and seven samples of defensive secretion (including six paired samples, i.e. from the same individual). One haemolymph sample was excluded since the colchicum alkaloid peaks were too small for automatic integration. Before HPLC-analysis, filled capillaries were stored at −80°C until extraction. To analyze alkaloids, whole capillaries including liquid samples were homogenized and extracted with methanol as described in the HPLC-methods section of the main manuscript. Statistical comparison of colchicum alkaloid concentrations between *S. saxatilis* defensive secretion and haemolymph was carried out using matched pair analysis based on six paired samples (i.e., six samples of haemolymph and six samples of secretions from the same individuals).

#### HPLC analysis of museum specimens

To test for the occurrence of colchicum alkaloids and cardiac glycosides in museum specimens of *S. saxatilis*, we incubated dry insects in 1 ml of methanol for at least one day and collected supernatants in fresh vials. After repeating this procedure twice (i.e., 3 ml methanol in total), pooled supernatants were evaporated under N2 and dissolved in 100 μl of methanol using a FastPrep homogenizer (6.5 m/s, 45 sec.). Before analyzing samples for colchicum alkaloids or cardenolides via HPLC using the respective methods (see above), samples were centrifuged (16,100 x g, 3 min) and filtered as described above.

#### Preparing seeds for HPLC

To compare toxins from host plant seeds to toxins sequestered by the bugs, seeds of *A. vernalis* (10.51 – 14.47 mg, commercial source), *C. autumnale* (5.91 – 10.03 mg, Berghausen, Germany, 2016), *D. purpurea* (59.44 – 63.72 mg, Eberbach, Germany, 2017), *E. crepidifolium* (10.19 – 10.65 mg, Schloßböckelheim, Germany, 2018), and *U. maritima* (2.89 – 7.66 mg, Aracena, Spain, 2016) were weighed and extracted with 0.5 ml (*A, vernalis, C. autumnale, U. maritima*) or 1 ml (*D. purpurea, E. crepidifolium*) methanol containing 0.01 mg/ml digitoxin (*A. vernalis*) or oleandrin (*D. purpurea*) as an internal standard (no internal standard for *C. autumnale, E. crepidifolium*, and *U. maritima*). For homogenization in a FastPrep homogenizer (2 x 6.5 m/s, 45 s), we used either zirconia beads for *D. purpurea* and *E. crepidifolium* or lysing matrix A (MP Biomedicals) for *A. vernalis*, *C. autumnale* and *U. maritima*. After grinding, samples were centrifuged (16,100 x g, 3 min) and supernatants were transferred to fresh vials. This procedure was repeated once for *D. purpurea* and *E. crepidifolium* seeds and twice for *A. vernalis*, *C. autumnale*, and *U. maritima* seeds. Supernatants of individual samples were pooled and evaporated under N2. Before HPLC-analyses, dried residues were dissolved in 500 (*C. autumnale, U. maritima*), 200 (*D. purpurea, E. crepidifolium*) or 100 μl methanol (*A. vernalis*) and analyzed as described above.

#### Evaluation of chromatograms

For chromatograms obtained from extracts of field-collected *H. superbus* from *D. purpurea, L. equestris, S. pandurus*, and *S. saxatilis* we considered all peaks of a sample showing compound specific absorption spectra (i.e. cardenolides, bufadienolides, and colchicum alkaloids) unless peaks were not automatically recognized by the software or had a signal to noise ratio < 2:1. For eggs of *L. equestris* and *S. saxatilis* as well as for the comparison between *S. saxatilis* secretion and haemolymph, the same approach was used. For samples obtained during seed mixture assays, we followed the procedure described above for *S. saxatilis*. Peaks in chromatograms from extracts of *H. superbus* and *L. equestris* were only taken into account if they were present in at least 70% or 60% of samples, respectively. The same approach was used for *H. superbus, L. equestris* larvae obtained from the feeding experiments with lacewings. For larvae of *S. saxatilis* and *S. pandurus* peaks were included that were present in at least 80% or 70% of samples, respectively. To evaluate field-collected *H. superbus* from *E. crepidifolium* we included all peaks that occurred in at least in 65% of samples. All datasets that were compared statistically were evaluated based on identical criteria.

#### Structural identification via reference compounds and liquid chromatography – mass spectrometry

To verify structural identity of selected cardenolides and colchicum alkaloids we compared HPLC retention times of chromatographic peaks obtained from field-collected *L. equestris* samples with authentic standards of k-strophanthoside (Roth, Germany), strophanthidin (PhytoLab, Germany) and cymarin (PhytoLab, Germany), cardenolides which are known to occur in *A. vernalis^47^*. For the same purpose, extracts from *H. superbus* were compared to authentic digitoxigenin (Sigma-Aldrich, Germany), digitoxin (Sigma-Aldrich, Germany), digoxigenin (Sigma-Aldrich, Germany), digoxin (Sigma-Aldrich, Germany), gitoxigenin (Santa Cruz Biotechnology, USA), gitoxin (EDQM, France), lanatoside C (Sigma-Aldrich, Germany), purpurea glycoside A (EDQM, France), and purpurea glycoside B (EDQM, France) which are known to occur in *D. purpurea*^75^, and erysimoside (Latoxan, France), known to occur in *E. crepidifolium*^76^. Similarly, we screened extracts of *S. pandurus* collected from *U. maritima* for the *Urginea* bufadienolides proscillaridin A (PhytoLab, Germany) and scillaren A (Sigma-Aldrich, Germany)^77^(Supplemental Figure 2e). Last, we compared chromatograms from field-collected *S. saxatilis* to the authentic colchicum alkaloids colchicoside, 2-demethylcolchicine, 3-demethylcolchicine (Toronto Research Chemicals, Canada), and colchicine (Roth, Germany) (Supplemental Figure 2e). In addition to the comparison of HPLC retention times, we compared mass spectra of the colchicum alkaloid compounds in the bug extracts to spectra of authentic alkaloid standards. Mass spectral analyses were performed on a Bruker micrOTOF-Q II mass spectrometer equipped with a Dionex Ultimate 3000 UHPLC and a C18 HPLC column (Kinetex C18, 2.6μ, 100A, 150 x 2.1 mm; Phenomenex). Compounds were separated by gradient elution with a flow rate of 150 mL min^-1^ and the following gradient of solvent A (0.1% formic acid in water) and solvent B (0.1% formic acid in acetonitrile): 0 to 2 min 16% B, 25 min 70% B, 30 min 95%. The column was eluted with 95% B solvent for an additional 10 min and re-equilibrated at the starting condition for 5 min. Sodium formate infusions at the beginning of each sample run were used for the mass calibration.

#### Effect of sequestered toxins on consumption of milkweed bug larvae by lacewing larvae

Milkweed bug and lacewing larvae were weighed before the trial and after exposition of bugs to lacewing larvae (after 12 hours for *H. superbus* and 8 hours for the other species) to assess consumption of milkweed bug larvae by lacewings. For statistical analysis, only dead bugs and corresponding lacewing larvae were included to ensure that milkweed bug larvae had been actually attacked by the lacewings. The aim of this experiment was to evaluate potential deterrence of sequestered plant compounds on lacewing feeding.

To assess the effect of sequestered toxins on remaining body mass of milkweed bug larvae after partial consumption by lacewings, remaining body mass was log_10_-transformed. We tested potential differences between dietary treatments using ANCOVA in JMP and included the initial mass of the intact milkweed bug larvae (i.e. before the experiment) as well as the initial mass of the lacewing larvae (as an estimate for body size) in our model. Furthermore, we tested for an interaction between the dietary treatment and the initial mass of the bugs. Since we carried out two experimental rounds for *L. equestris*, we included ‘experiment’ as a blocking term. For the remains of one out of 52 *L. equestris* individuals and four out of 40 *S. saxatilis* individuals, body mass was zero (i.e. the mass was below the detection limit of the balance used), thus these data were removed before log_10_-transformation.

For the comparison of lacewing weights after feeding on milkweed bug larvae, we used the same model as for milkweed bugs using untransformed data with final mass of the lacewings as the main effect. We included the initial weight of bugs and lacewings as model effects. Due to the two experimental rounds for *L. equestris*, we again included ‘experiment’ as a blocking term. Initial masses of milkweed bugs and lacewings across treatments (i.e. bugs raised on sunflower vs. toxic seeds) were compared by two-tailed Student’s t-tests in JMP except of the experiment with *L. equestris* that we analyzed with an ANCOVA again using ‘experiment’ as a blocking term due to the two experimental rounds for this species.

#### Hand rearing juvenile birds for predation assays

Juvenile great tits were obtained from a population breeding in nest-boxes in mixed woods at the outskirts of Prague. The juveniles were taken from nest-boxes when 12–15 days old, and hand reared in the laboratory. This way, they were naive in regard to experience with any kind of unpalatable or warningly colored prey. The birds were kept in artificial nests until fledging and then housed in groups of three or four in indoor cages (60 x 50 x 50cm) under illumination and temperature regimes simulating natural conditions. Their diet consisted of mealworms (*Tenebrio molitor* larvae) and commercial mixtures for hand-rearing passerine birds (Handmix, NutriBird, Gold Patee and Uni Patee Premium, Orlux). Birds were used for predations assays when at least 35-days old and fully independent. We ringed the birds individually and released them back to the locality of capture within a few days after experimentation. For experiments with great tits, we obtained permissions from the Environmental Department of Municipality of Prague (S-MHMP-83637/2014/OZP-VII-3/R-8/F), Ministry of Agriculture (13060/2014-MZE-17214), and Ministry of the Environment of the Czech Republic (42521/ENV/14-2268/630/14).

#### Attack latencies and duration of discomfort-indicating behavior in avian predators

In each trial of behavioral assays with great tits as predators, we recorded latency of the first attack and duration of discomfort-indicating behavior observed in birds (beak wiping and head shaking). Data were log_10_-transformed to meet the assumptions of normality and homogeneous variance and analyzed in the lme4^78^ and geepack^54^ packages in R^53^.

Attack latencies in the first trial were analyzed using a general linear model (ANOVA) with bug species and host plant toxicity entered as fixed effects. Latencies of the first attacks were also compared between bugs and control crickets by paired t-test. Changes in attack latencies over the three trials were analyzed using a generalized estimating equation model (GEE) in a subset of data including only the birds that attacked all three bugs offered. Trial number, bug species and host plant toxicity were entered as fixed effects and bird individual (id) as a random effect.

Durations of discomfort-indicating behavior recorded during the first trial were analyzed using a general linear model (ANOVA) with bug species and host plant toxicity entered as fixed effects. To evaluate whether the general effect of host plant toxicity on discomfort-indicating reactions of birds also holds for each of the milkweed bug species studied, similar models were run separately for each bug species.

#### Survival of milkweed bugs compared to control palatable prey

We used GEE models (package geepack^54^ in R^53^) to compare survival rates of milkweed bugs raised on sunflower with survival rates of control crickets. The data included only the birds that attacked all three bugs offered and were analyzed for each milkweed bug species separately. Prey (cricket versus bug) and trial number were entered as fixed effects and bird individual (id) as a random effect.

#### Analysis of bugs eaten by avian predators

Following the observations of birds consuming the bugs in particular trials, we examined the remaining parts of the bugs using a stereomicroscope to determine what body parts the birds were able to consume. In a subset of data from first trials including only the cases when the bugs were killed, we analyzed an effect of host plant on the probability that at least part of the bug would be consumed. The data were analyzed separately for each species by generalized linear models (GLM) with binomial errors using the lme4 package^78^ in R^53^, and the hostplant was entered as a fixed effect. In addition, we compared frequency of different body parts of bugs consumed by the birds using Fisher’s exact test.

### Supplementary Results

#### Developmental time on different diets

*H. superbus* developed equally fast on all diets (F_3,35_ = 2.22, p = 0.103, n = 9 for the seed mixture without *Digitalis* and n = 10 for all other treatments) while the dietary treatment affected developmental time in *L. equestris* (F_3,40_ = 7.168, p < 0.001, n = 11 for all diets). Larvae needed longer to reach the adult stage on *A. vernalis* seeds compared to sunflower seeds and the seed mixture containing *A. vernalis* seeds (LSMeans Tukey HSD: p = 0.002; p = 0.001) but not compared to the *A. vernalis-free* seed mixture (LSMeans Tukey HSD: p = 0.097). Growth between the other diets was not different (LSMeans Tukey HSD: seed mixture vs. seed mixture with *A. vernalis* seeds, p = 0.3; seed mixture vs. sunflower, p = 0.48; sunflower vs. seed mixture with *A. vernalis*, p = 0.987).

#### Sequestration on toxic seeds across different diets

On seed mixtures containing low amounts of toxic seeds of either *Digitalis, Adonis*, or *Colchicum*, sequestration of plant toxins was reduced by approximately 50 % in *H. superbus* (Welch’s test: F_1,19_ = 18.863, p < 0.001) and *S. saxatilis* (Welch’s test: F_3,20_ = 83.568, p < 0.001; Games-Howell post-hoc test significant at p < 0.05 for all comparisons except of pure sunflower seeds and the seed mixture without *C. autumnale* seeds) while *L. equestris* accumulated only about 10 % of the amount of cardenolides it contained on the pure toxic diet (Welch’s test: F_1,10_ = 66.206, p < 0.001, (Figure 2a-c). Notably, we found toxins in all (n = 32) specimens sampled from the seed mixture treatments including *D. purpurea*, *A. vernalis*, or *C. autumnale* seeds, showing that toxic seeds were always accessed by the bugs, even if only available in small numbers within mixtures.

#### Natural history remarks

When collecting milkweed bugs in the field we also observed feeding behavior and host plant use. In general, feeding was mainly restricted to fruits or flowers. *H. superbus* was only observed on *D. purpurea* plants and substrates such as bark and stumps but never on other plants. Early in the season, we found the bugs walking on *Digitalis*-leaves and stems or feeding on flowers. Later in the season, we typically found *H. superbus* in the opened ripe *Digitalis*-pods (adults and larvae). In a *Digitalis-free* habitat, we found *H. superbus* exclusively on seedpods of *E. crepidifolium*. In this habitat, the insects may also suck on the fleshy leaves of *Sedum* spec., a cushion plant that they may use as a refuge. The feeding ecology of *L. equestris* has been described elsewhere^45^. Our own observations confirm that in *A. vernalis* habitats, *Adonis* is the primary host plant early in the season. We and others recorded *S. saxatilis* feeding on more than 40 plant species from more than 15 botanical families (Supplemental Table 3). *S. saxatilis* oviposits into *C. autumnale* seedpods and early larval stages of *S. saxatilis* were only observed in ripe fruits of *C. autumnale* (Supplemental Figure 6).

#### Effects of sequestered Digitalis-cardenolides on consumption of H. superbus by lacewing larvae

Although individuals of *H. superbus* raised on sunflower seeds had a greater body mass compared to individuals raised on *Digitalis* seeds before the experiment (t = 2.822, df = 21, p = 0.01, n = 11 for sunflower-raised and n = 12 for *D. purpurea-raised* bugs), carcasses from sunflower-raised bugs were lighter after attack compared to carcasses from *Digitalis* raised bugs (F_1,18_ = 39.205, p < 0.001, Supplemental Figure 7a). The body mass of bugs prior to lacewing feeding (‘initial mass’) affected the remaining mass (F_1,18_ = 23.825, p < 0.001), while initial mass of lacewings (i.e. before feeding on a milkweed bug larva) had no effect (F_1,18_ = 2.636, p = 0.122). There was no interaction between initial mass of bugs and treatment (F_1,18_ = 0.269, p = 0.611) on remaining mass. In accordance with lower consumption of *Digitalis*-raised bugs, lacewing larvae gained more body mass when feeding on sunflower raised-bugs compared to feeding on *Digitalis*-raised bugs (F_1,18_ = 48.833, p < 0.001, Supplemental Figure 7e). The initial masses of bugs and lacewings influenced the final lacewing mass (F_1,18_ = 9.091, p = 0.007; F_1,18_ = 306.9, p < 0.001). In addition, final mass of lacewings was affected by an interaction between the initial mass of bugs and their diet (F_1,18_ = 7.258, p = 0.015). The initial mass of lacewing larvae before the trial was not different (t = −0.059, df = 21, p = 0.953).

#### Effects of sequestered Adonis-cardenolides on consumption of L. equestris by lacewing larvae

The initial mass of bugs (analyzed by ANCOVA to account for experimental blocks, n = 21 for sunflower-raised and n = 30 for *A. vernalis-raised* bugs) neither differed between diets (F_1,48_ = 1.026, p = 0.316) nor between the two experimental rounds (F_1,48_ = 0.046, p = 0.831). Consumption of milkweed bugs by lacewing larvae was higher on sunflower-raised bugs compared to bugs raised on *Adonis* (F_1,45_ = 9.170, p = 0.004, Supplemental Figure 7b). Again, the remaining mass of bugs was affected by their inital mass (F_1,45_ = 23.854, p < 0.001) and there was no interaction between initial mass and treatment (F_1,45_ = 0.235, p = 0.630). The mass of lacewing larvae before the trial was not different across diets and experiments (F_1,49_ = 0.087, p = 0.77; F_1,49_ = 0.953, p = 0.334; n = 22 for lacewings feeding on sunflower and n = 30 for lacewings feeding on *A. vernalis-raised* bugs) but predicted how much body mass of the bugs was consumed (F_1,49_ = 6.286, p = 0.016). In accordance with the observed loss of body mass in the bugs, lacewings gained more body mass when feeding on sunflower-raised bugs compared to *Adonis*-raised bugs (F_1,46_ = 8.02, p = 0.007, Supplemental Figure 7f). The initial masses of bugs and lacewings determined the final body mass of the lacewing larvae (F_1,46_ = 6.286, p = 0.016; F_1,46_ = 98.245, p < 0.001). While the interaction between diet and initial mass of the bugs had an effect on lacewing final mass the round of experiment had no influence (F_1,46_ = 4.3, p = 0.044; F_1,46_ = 0.045, p = 0.833).

#### Effects of sequestered Urginea-bufadienolides on consumption of S. pandurus by lacewing larvae

Initial mass of bugs did not differ between the two diets (sunflower vs. *Urginea* seeds, t = −0.173, df = 36, p = 0.864, n = 19 each). In contrast to the other species, we only found a marginal effect of the diet on remaining milkweed bug mass after the lacewing attack (F_1,33_ = 3.79, p = 0.06, Supplemental Figure 7c). The initial mass of the bugs affected the remaining mass after the experiment (F_1,33_ = 21.202, p < 0.001) and there was a significant interaction between diet and initial mass affecting remaining body mass (F_1,33_ = 5.602, p = 0.024). The initial mass of lacewing larvae had no effect on the remaining mass of the bugs after consumption (F_1,33_ = 1.199, p = 0.282). Nevertheless, lacewing larvae were heavier after feeding on sunflower raised *S. pandurus* larvae compared to feeding on bugs raised on *Urginea* seeds (F_1,33_ = 195.969, p < 0.001, Supplemental Figure 7g). Besides the diet, the initial mass of bugs and lacewings as well as the interaction between initial mass of bugs and diet affected body mass of lacewings after the trial (F_1,33_ = 44.982, p < 0.001; F_1,33_ = 174.744, p < 0.001; F_1,33_ = 20.639, p < 0.001). The mass of lacewing larvae was not different across treatments before the experiment (t = 1.052, df = 32, p = 0.301).

#### Effects of sequestered Colchicum-alkaloids on consumption of S. saxatilis by lacewing larvae

Initial mass was higher in *Oncopeltus* (n = 16) than in *S. saxatilis* (n = 20; t = 12.533, df = 30, p < 0.001, Supplemental Figure 7d) but after lacewing predation, the pattern was reversed (F_1,31_ = 57.516, p < 0.001). The initial masses of bugs and lacewings did not affect remaining mass of bugs after attack (F_1,31_ = 2.91, p = 0.098 and F_1,31_ = 0.012, p = 0.914) and there was no interaction between the initial mass of bugs and treatment affecting remaining body mass after attack (F_1,31_ = 0.143, p = 0.708). Lacewing larvae assigned to *O. fasciatus* (n = 20) larvae were heavier compared to lacewing larvae assigned to larvae of *S. saxatilis* (n = 20; t = 4.324, df = 38, p < 0.001). Nevertheless, this difference was more pronounced after the trial (2-fold vs. 1.75-fold, F_1,35_ = 129.881, p < 0.001, Supplemental Figure 7h). The initial masses of bugs and lacewings affected the final mass of lacewings (F_1,35_ = 12.225, p = 0.001; F_1,35_ = 801.952, p < 0.001) and there was an interaction between the initial mass of bugs and treatment affecting lacewing final mass (F_1,35_ = 41.971, p < 0.001).

#### Attack latency and duration of discomfort-indicating behavior in avian predators

In first encounters with novel prey, birds did not hesitate longer before attacking milkweed bugs than before attacking control crickets (paired t-test, t = 0.173, df = 99, p = 0.863; Supplemental Figure 8a). Furthermore, attack latencies were influenced neither by milkweed-bug species (ANOVA, F_2,94_ = 1.241, p = 0.294) nor by host plant toxicity (ANOVA, F_1,94_ = 1.876, p = 0.117). Thus, we have not found any evidence of an innate bias against warningly coloured milkweed bugs in juvenile great tits or for before-attack defensive effects of sequestered host plant toxins. Nevertheless, birds that attacked all three bugs in a row hesitated longer before attacking the bugs raised on toxic host plants upon repeated encounters (GEE, trial, χ^2^_1_ = 0.931, p = 0.335; host plant, χ^2^_1_ = 27.935, p < 0.001; trial: host plant interaction, χ^2^_1_ = 12.384, p < 0.001; Supplemental Figure 8b).

When handling the bugs, birds often responded by discomfort-indicating behavior (head shaking and beak wiping). Duration of this behavior in the first trial was similar across all bug species tested (ANOVA, F_2,94_ = 1.928, p = 0.151). However, the birds tested with bugs raised on toxic host plants spent more time by discomfort–indicating behavior than the birds tested with bugs from control sunflower (ANOVA, F_1,94_ = 37.980, p < 0.001; Supplemental Figure 9). There was no interaction between the two factors (ANOVA, F_2,94_ = 0.987, p < 0.001 = 0.376). When analyzed separately for each species, birds spent longer time by discomfort-indicating behavior when attacking and handling bugs raised on toxic host plants than when handling bugs from sunflower (ANOVA, *S. saxatilis:* F_1,38_ = 11.271, p = 0.002, *L. equestris:* F_1,38_ = 20.962, p < 0.001, *H. superbus:* F_1,18_ = 6.177, p = 0.023; Supplemental Figure 9a-c). This result indicates that chemicals sequestered from host plants cause stronger aversion in avian predators upon direct contact than the secretion of metathoracic glands alone.

#### Survival of milkweed bugs compared to control palatable prey

In all three species of milkweed bugs, the probability to survive repeated attacks by avian predators was higher for the sunflower-raised bugs than for control crickets (GEE, *S. saxatilis:* χ^2^_1_ = 6.235, p = 0.012; *L. equestris:* χ^2^_1_ = 6.534, p = 0.011; *H. superbus:* χ^2^_1_ = 3.899, p = 0.048). These results indicate that the line of defence based on the secretion of metathoracic scent glands is effective by itself, even though its sole effect is considerably smaller compared to when it is combined with sequestration of host plant chemicals.

#### Consumption of bugs by avian predators

In all three milkweed bug species tested, host plant affected the probability that the birds would eat at least a part of the bug attacked and killed in the first trial (GLM, *S. saxatilis:* χ^2^_1_,24 = 13.214, *P* < 0.001; *L. equestris:* χ^2^_1_,27 = 5.849, *P* = 0.016; *H. superbus:* χ^2^_1_,15 = 7.945, *P* = 0.005). Whereas the birds frequently ate at least some parts of sunflower-raised bugs, consumption of bugs raised on toxic host plants was exceptionally rare. Out of 20 birds tested with *S. saxatilis* and *L. equestris*, only two and four birds, respectively, ate some parts of *Colchicum*- and *Adonis*-raised bugs, and in all cases it was only a small part of the abdomen (fat body). Likewise, out of 10 birds tested with *H. superbus*, only one bird consumed a small part of the abdomen of a *Digitalis*-raised bug. Some of the birds tested with bugs from non-toxic hostplants consumed whole bugs or left only few fragments of cuticle, but they nevertheless consumed parts of the abdomen (usually the fat body) significantly more frequently than other parts of the bug (two tailed Fisher’s exact test, *S. saxatilis: P* < 0.001; *L. equestris: P* < 0.001; *H. superbus: P* < 0.021).

#### Structural identification of sequestered compounds and comparison to seed extracts

We were not able to identify individual compounds by comparing insect extracts to nine cardenolide reference compounds known to occur in *D. purpurea*^75^ (Supplemental Figure 2a). Similarly, the cardenolide profile from *D. purpurea* seeds revealed only little putative overlap with cardenolides sequestered by the insects (Supplemental Figure 3a) indicating extensive metabolic transformation by the insect. In contrast, comparison of *H. superbus* collected from *E. crepidifolium* revealed clear overlap with seed extracts (Supplemental Figure 3d) and we identified one substance as erysimoside based on the retention time of a commercial reference compound (Supplemental Figure 2d). The comparison of seed extracts from *A. vernalis* to extracts of *L. equestris* revealed no clear similarity (Supplemental Figure 3b). Three cardenolides sequestered by *L. equestris* were putatively identified as k-strophanthosid, strophanthidin and cymarin based on authentic reference standards (Supplemental Figure 2b). The comparison of the HPLC profiles between insect extracts and *Urginea* seeds suggested that at least the dominant bufadienolides found in the insect are identical to compounds found in the seeds (Supplemental Figure 3c). Nevertheless, we did not detect the bufadienolides scillaren A and proscillaridin A (Supplemental Figure 2e) that are reported to occur in bulbs of *U. maritima*^77^.

In *S. saxatilis* from Berghausen we identified 12 peaks with absorption spectra similar to colchicine. Three of the four dominant peaks present in all extracts and accounting for > 90 % of the observed alkaloids were identified as colchicine (34.6 %), 2-demethyl-colchicine (16.2 %), and 3-demethyl-colchicine (20.1 %) based on retention time and molecular mass using authentic standards (Supplemental Figure 2c). We furthermore identified the colchicine glycoside colchicoside as a minor compound. In specimens obtained from Nüstenbach, we detected only nine peaks eight of which were identical with the ones observed in *S. saxatilis* from Berghausen including colchicine, 2 and 3-demethyl-colchicine, and colchicoside. Seeds of *C. autumnale* contained colchicine and colchicoside but none of the related alkaloids found in the bugs (Supplemental Figure 3e).

#### Injection experiments with ouabain and colchicine

*P. apterus* showed clear signs of intoxication in a dose-dependent manner for both toxins, ouabain (Cochrane-Armitage trend test, Z = −4.006, p < 0.001) and colchicine (Cochrane-Armitage trend test, Z = −4.887, p < 0.001, Supplemental Figure 10a,d). While unaffected by ouabain (Figure 4), *O. fasciatus* responded to colchicine in a dose-dependent manner (Cochrane-Armitage trend test, Z = −3.453, p < 0.001, Supplemental Figure 10b). At a dose of 5 μg per animal, 100% of individuals were affected while *S. saxatilis* tolerated up to 30 μg which was the highest dose tested (p = 0.467 compared to specimens injected with 5 μg ouabain that were all surviving; two tailed Fisher’s exact test, Supplemental Figure 10e). *S. pandurus*, although being a congener of *S. saxatilis*, was unable to tolerate colchicine and was affected in a dose dependent fashion (Cochrane-Armitage trend test, Z = −3.933, p < 0.001, Supplemental Figure 10c).

---

**Supplemental Table 1.** Plant species used for seed mixture experiments. Botanical families in parentheses. For *S. saxatilis* and *H. superbus* mean weights of seeds per plant species ranged from 12 - 13 mg (with 91 % of plant species being close to 12 mg) and from 9 - 13 mg for *L. equestris*, with 87 % of host seed species being close to 9 mg per Petri dish. Deviations from 9 mg are due to the extensive seed sizes of certain species (*T. pratensis*) i.e. individual seeds exceeded 9 mg.

| Plant species used for *S. saxatilis* | Plant species used for *H. superbus* | Plant species used for *L. equestris* |
| --- | --- | --- |
| *Achillea millefolium* (Asteraceae) | *Achillea millefolium* (Asteraceae) | *Achillea millefolium* (Asteraceae) |
| *Bupleurum falcatum* (Apiaceae) | *Bupleurum falcatum* (Apiaceae) | *Angelica archangelica* (Apiaceae) |
| *Centaurea jacea* (Asteraceae) | *Centaurea jacea* (Asteraceae) | *Centaurea scabiosa* (Asteraceae) |
| *Cichorium intybus* (Asteraceae) | *Cichorium intybus* (Asteraceae) | *Cichorium intybus* (Asteraceae) |
| *Daucus carota* (Apiaceae) | *Daucus carota* (Apiaceae) | *Cirsium arvense* (Asteraceae) |
| *Hieracium pilosella* (Asteraceae) | *Hieracium pilosella* (Asteraceae) | *Dactylis glomerata* (Poaceae) |
| *Origanum vulgare* (Lamiaceae) | *Origanum vulgare* (Lamiaceae) | *Daucus carota* (Apiaceae) |
| *Plantago major* (Plantaginaceae) | *Plantago major* (Plantaginaceae) | *Hieracium pilosella* (Asteraceae) |
| *Tanacetum vulgare* (Asteraceae) | *Tanacetum vulgare* (Asteraceae) | *Pimpinella saxifraga* (Apiaceae) |
| *Taraxacum officinale* (Asteraceae) | *Taraxacum officinale* (Asteraceae) | *Plantago major* (Plantaginaceae) |
| *Tragopogon pratensis* (Asteraceae) | *Tragopogon pratensis* (Asteraceae) | *Taraxacum officinale* (Asteraceae) |
|  |  | *Thymus serpyllum* (Lamiaceae) |
|  |  | *Tragopogon pratensis* (Asteraceae) |
|  |  | *Urtica dioica* (Urticaceae) |
|  |  | *Vincetoxicum hirundinaria* (Apocynaceae) |

**Supplemental Table 2.**
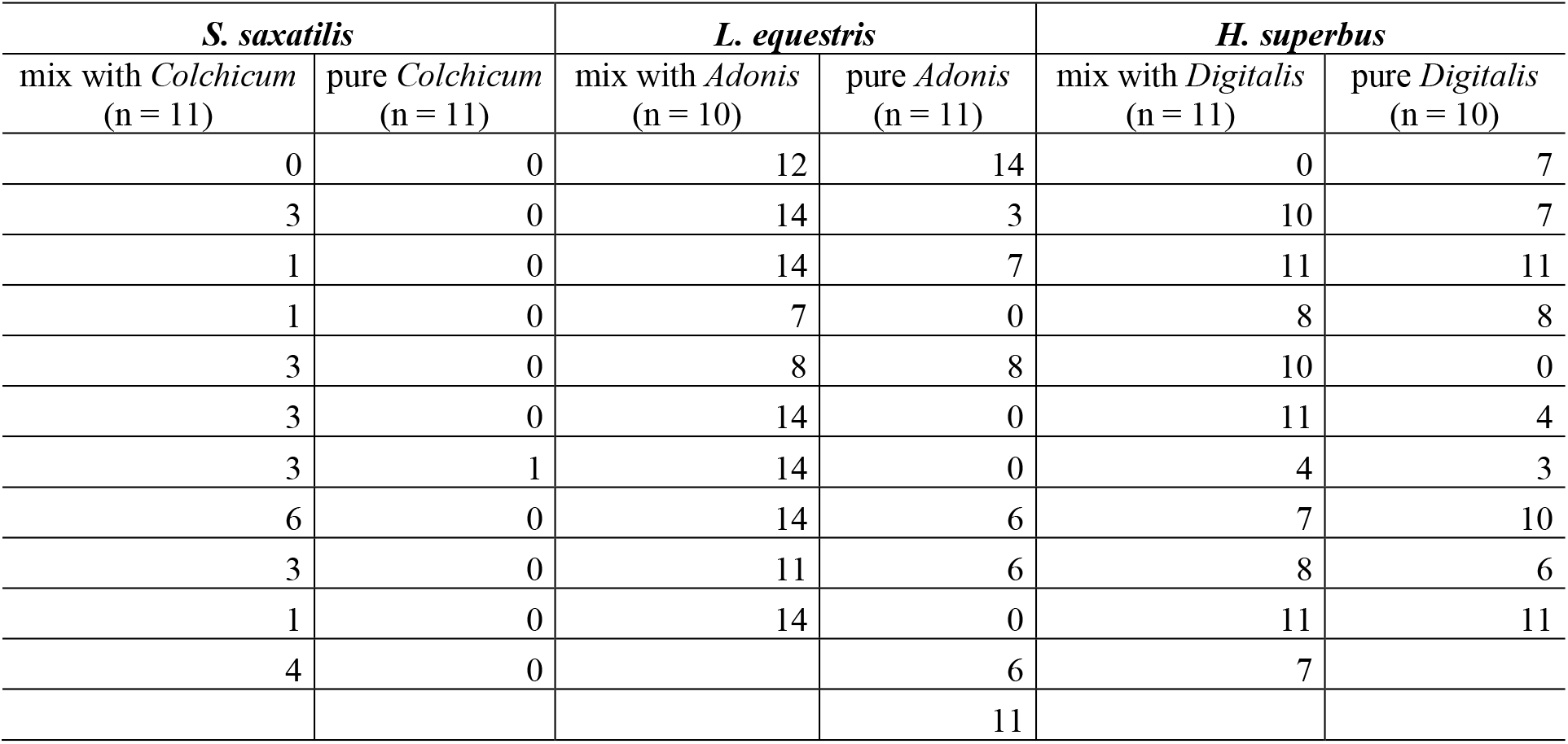
Number of days bugs were spending on sunflower seeds as adults (i.e. with a continuous gut lumen) for purging digestive tracts from potential residues of dietary toxins (cardenolides or colchicum alkaloids).

| *S. saxatilis* |  | *L. equestris* |  | *H. superbus* |  |
| --- | --- | --- | --- | --- | --- |
| mix with *Colchicum*<br>(n = 11) | pure *Colchicum*<br>(n = 11) | mix with *Adonis*<br>(n = 10) | pure *Adonis*<br>(n = 11) | mix with *Digitalis*<br>(n = 11) | pure *Digitalis*<br>(n = 10) |
| 0 | 0 | 12 | 14 | 0 | 7 |
| 3 | 0 | 14 | 3 | 10 | 7 |
| 1 | 0 | 14 | 7 | 11 | 11 |
| 1 | 0 | 7 | 0 | 8 | 8 |
| 3 | 0 | 8 | 8 | 10 | 0 |
| 3 | 0 | 14 | 0 | 11 | 4 |
| 3 | 1 | 14 | 0 | 4 | 3 |
| 6 | 0 | 14 | 6 | 7 | 10 |
| 3 | 0 | 11 | 6 | 8 | 6 |
| 1 | 0 | 14 | 0 | 11 | 11 |
| 4 | 0 |  | 6 | 7 |  |
|  |  |  | 11 |  |  |

**Supplemental Table 3.**
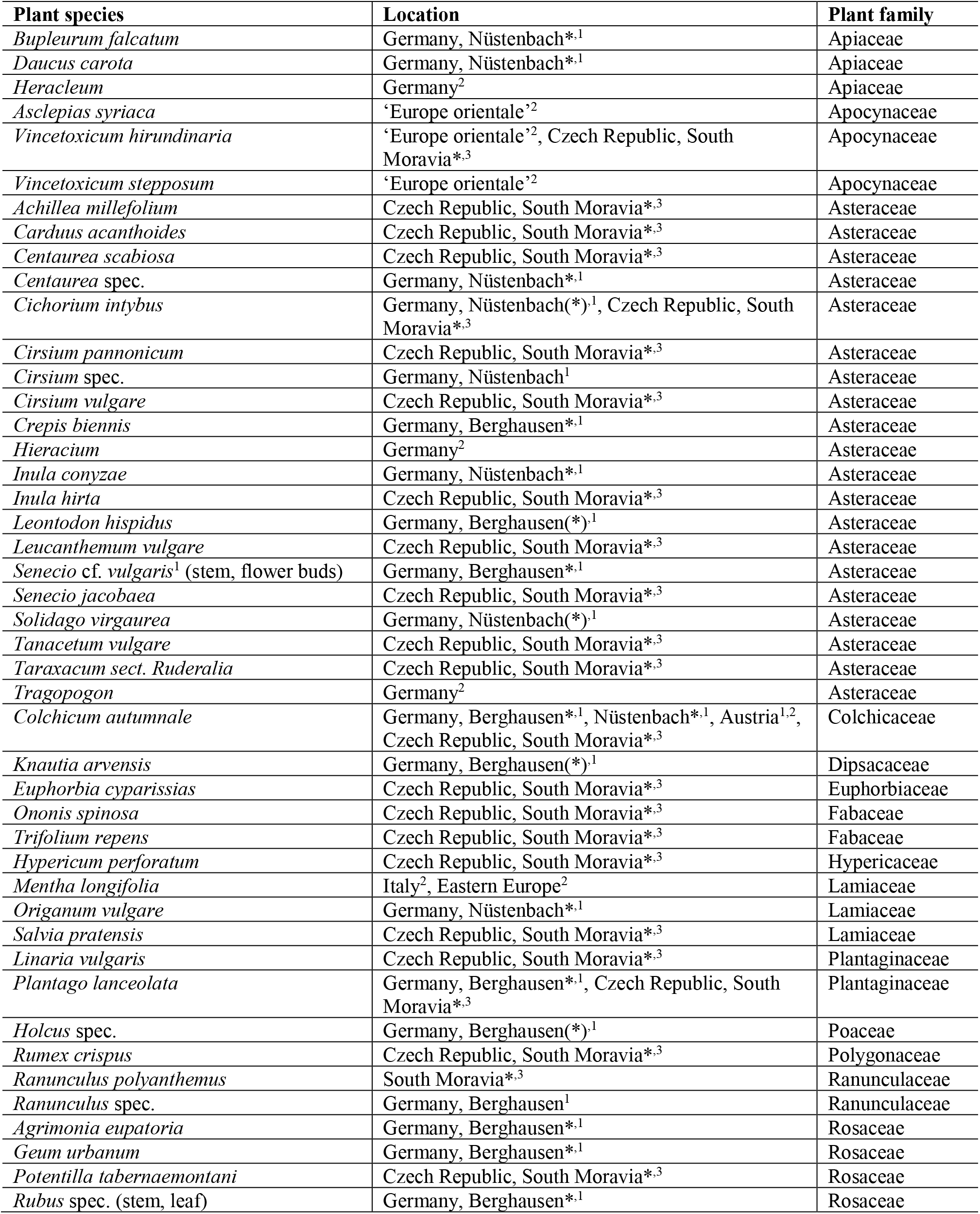
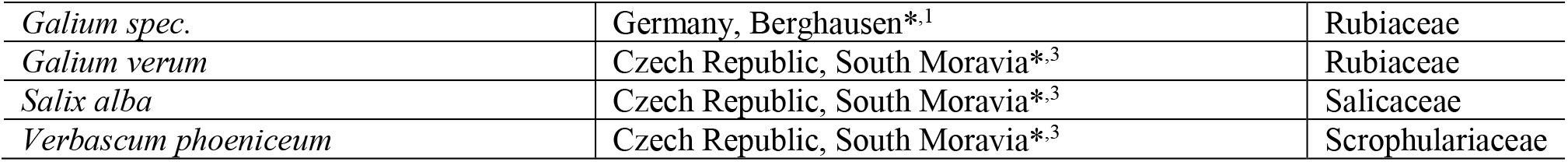
Host plants recorded for *S. saxatilis* in the field. Superscript numbers indicate sources of host plant data: ^1^ = own observation, ^2^ = Péricart (1998)^26^, ^3^ = Banar (2003)^79^. Asterisks indicate actual feeding observations.

**Supplemental Figure 1.**
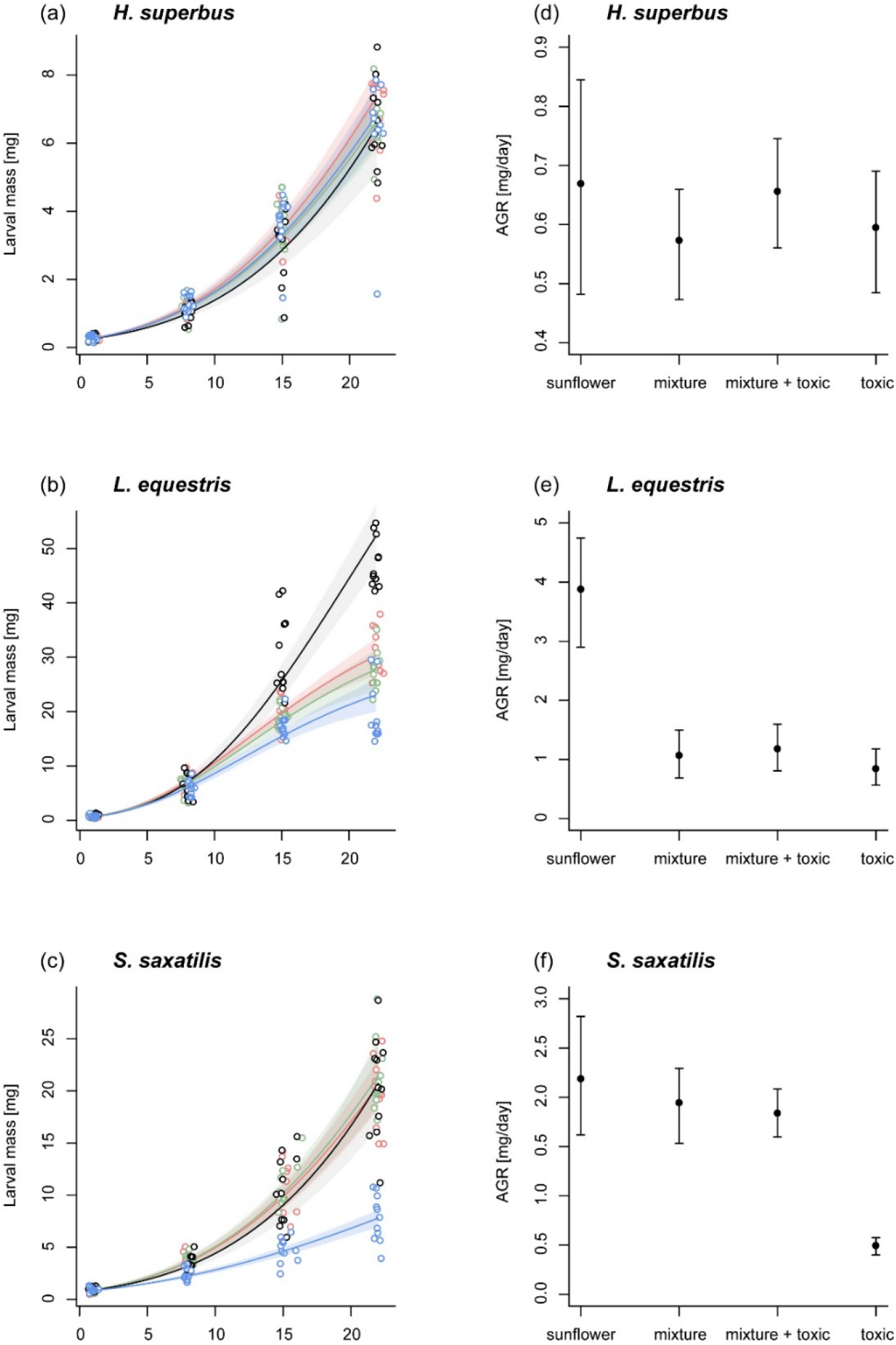
Growth of milkweed bug larvae on four different seed diets. (a-c) Weight gain of *H. superbus, L. equestris*, and *S. saxatilis* larvae feeding on sunflower seeds (control, black), a non-toxic seed mixture (green), a seed mixture with toxic seeds (red), or toxic seeds only (blue). Points are average weights of 1-3 larvae per petri dish (replicates), measured sequentially over three weeks. Growth was modelled as an asymptotic process using a non-linear mixed effects model (function *nlme* with *SSasymp* in R) with log-weight as the response, a random effect of petri dish to account for repeated measures, and fixed effects of species and treatment on individual model parameters (*K:* species × treatment, F_6,374_ = 165.4, p < 0.001; *M_0_*: species, F_2,374_ = 202.7, p < 0.001; *log-rate constant:* species, F_2,374_ = 83.7, p < 0.001, treatment, F_3,374_ = 4.9, p = 0.002). Solid lines are model predictions, and shaded areas are 95% population prediction intervals, generated by drawing random values from the estimated sampling distribution of each regression parameter. (d-e) Absolute growth rates for *H. superbus*, *L. equestris*, and *S. saxatilis* larvae feeding on different seed diets. AGR values were calculated for growth on day 22 (final day of the experiment), using the model parameters from the asymptotic regression model displayed in panels a-c. Error bars are 95% population prediction intervals.

**Supplemental Figure 2.**
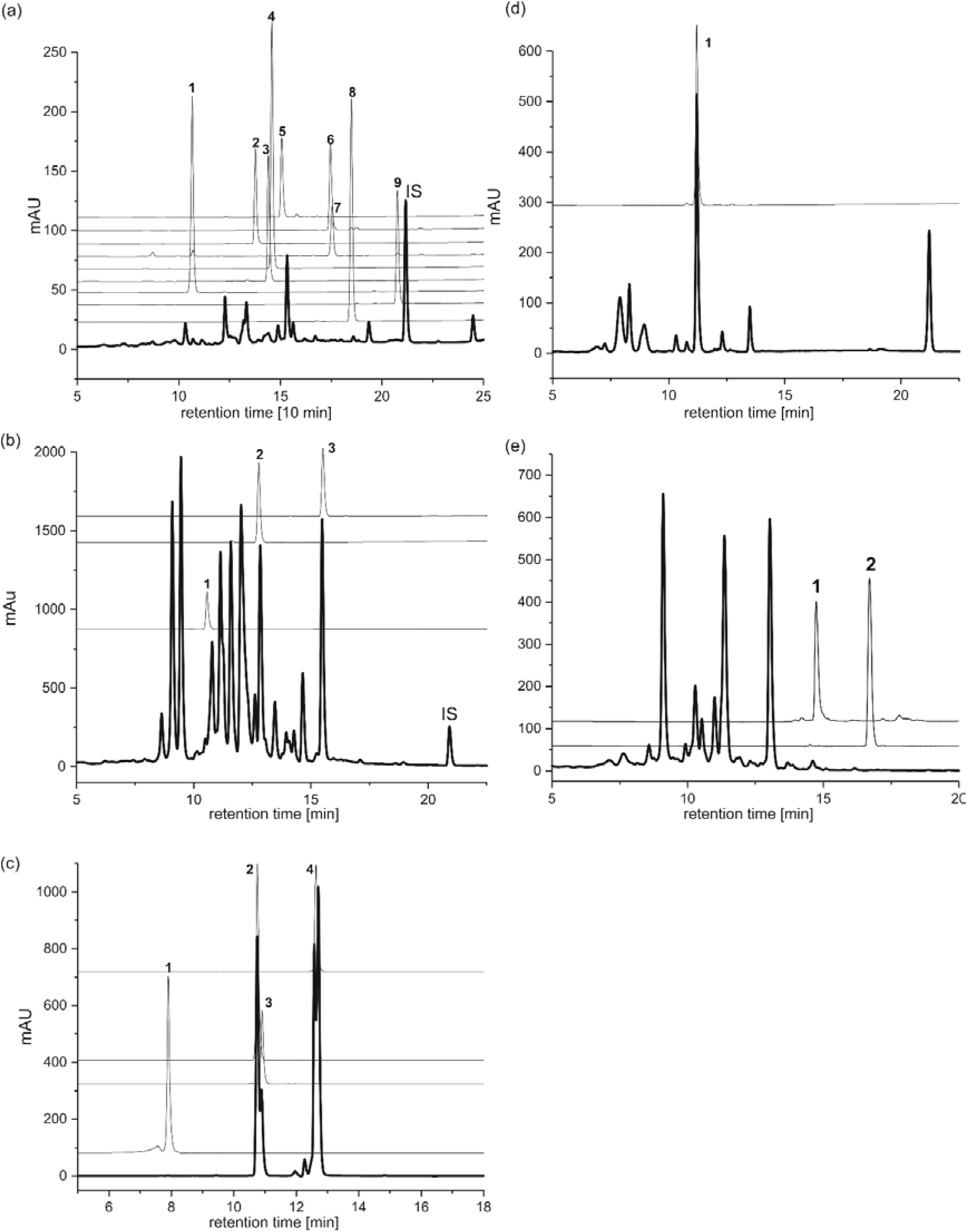
Comparison of HPLC-chromatograms obtained from field-collected milkweed bugs with authentic reference compounds. (a) *H. superbus* from *D. purpurea* compared to compounds reported from *D. purpurea*: 1 = digoxigenin, 2 = lanatoside C, 3 = digoxin, 4 = gitoxigenin, 5 = purpurea glycoside B, 6 = purpurea glycoside A, 7 = gitoxin, 8 = digitoxigenin, 9 = digitoxin. (b) *L. equestris* collected from *A. vernalis* compared to 1 = k-strophanthosid, 2 = strophanthidin and 3 = cymarin that all occur in *A. vernalis*. (c) *S. saxatilis* from *C. autumnale* compared to authentic colchicum alkaloid standards: 1 = colchicoside, 2 = 3-demethyl colchicine, 3 = 2-demethyl colchicine, and 4 = colchicine. (d) *H. superbus* from *E. crepidifolium* compared to erysimoside (1). (e) *S. pandurus* from *U. maritima* compared to the *Urginea* bufadienolides 1 = scillaren A, 2 = proscillaridin A.

**Supplemental Figure 3.**
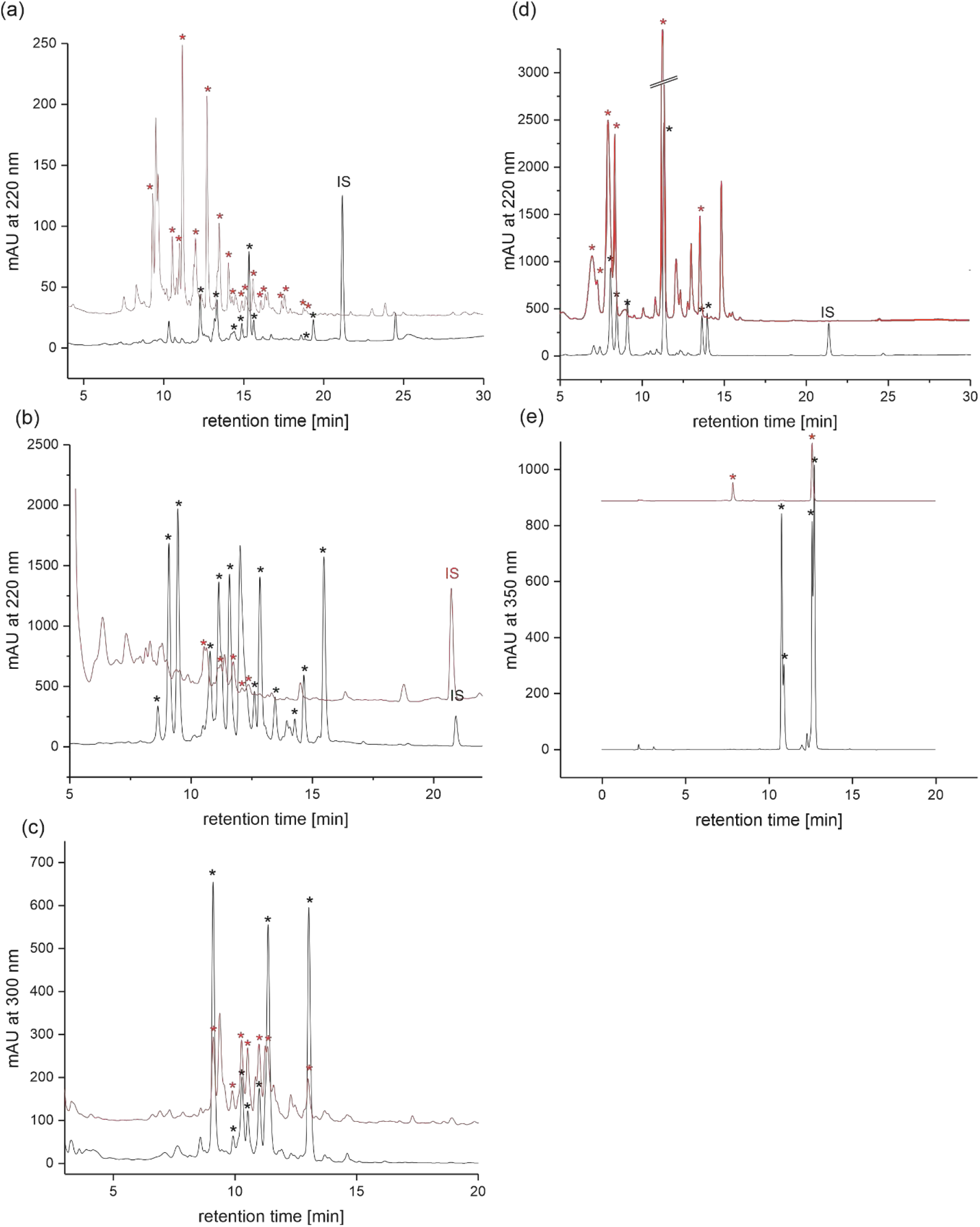
Comparison of milkweed bug and host plant seed extracts with HPLC-DAD. (a) *H. superbus* vs. *D. purpurea* seeds, (b) *L. equestris* vs. *A. vernalis* seeds, (c) *S. pandurus* vs. *U. maritima* seeds, (d) *H. superbus* vs. seeds of *E. crepidifolium*, (e) *S. saxatilis* vs. *C. autumnale* seeds. Top chromatograms (red) always represent seed; bottom chromatograms (black) always represent insect extracts. Asterisks indicate the most prominent cardiac glycoside or colchicum alkaloid peaks, respectively.

**Supplemental Figure 4.**
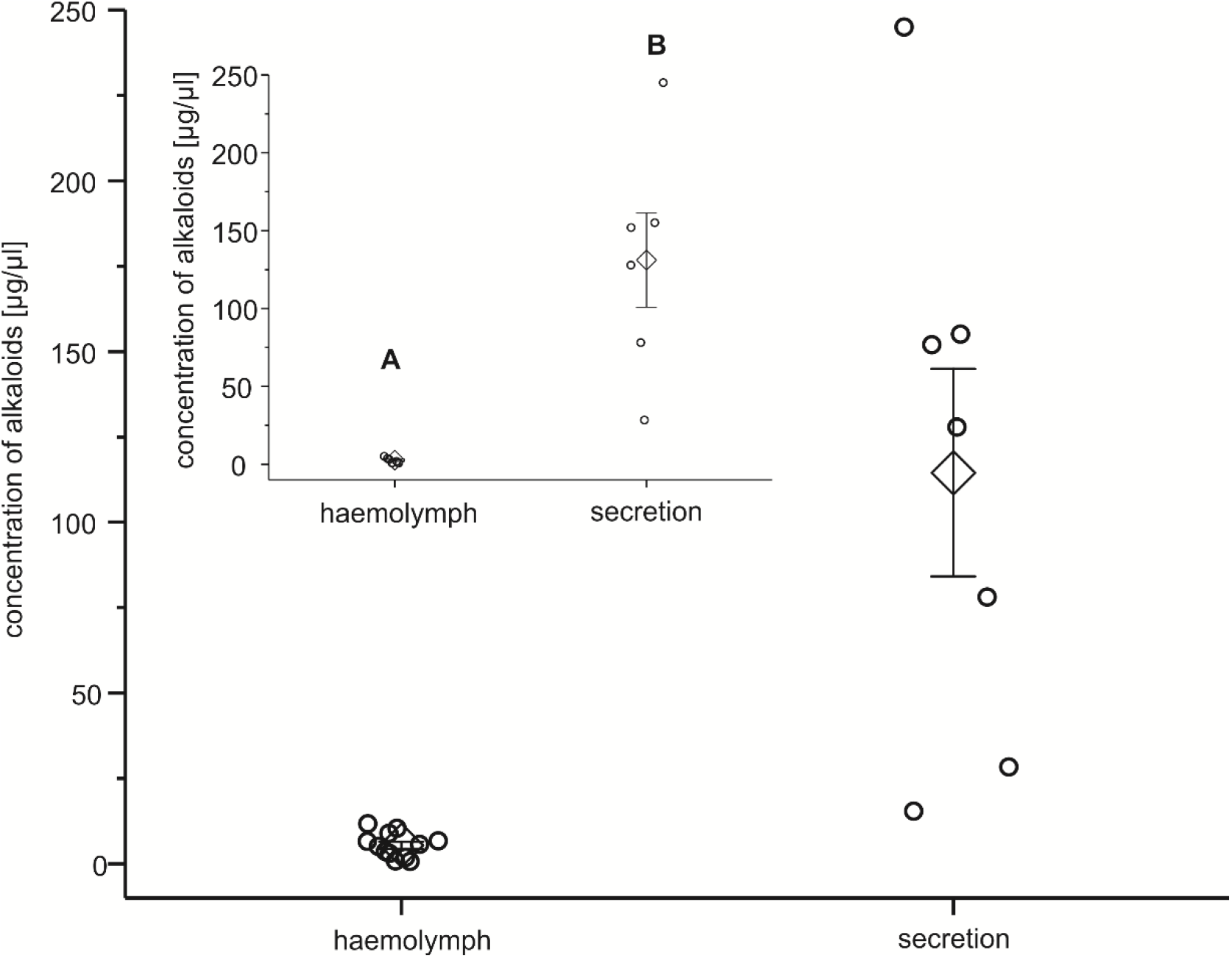
Concentration of colchicum alkaloids in *Spilostethus saxatilis* haemolymph (n = 12) and defensive secretion (n = 7). Diamonds are means ± SE, circles represent jittered raw data. Subset of six paired samples obtained from the same individuals used for statistical comparison (inset).

**Supplemental Figure 5.**
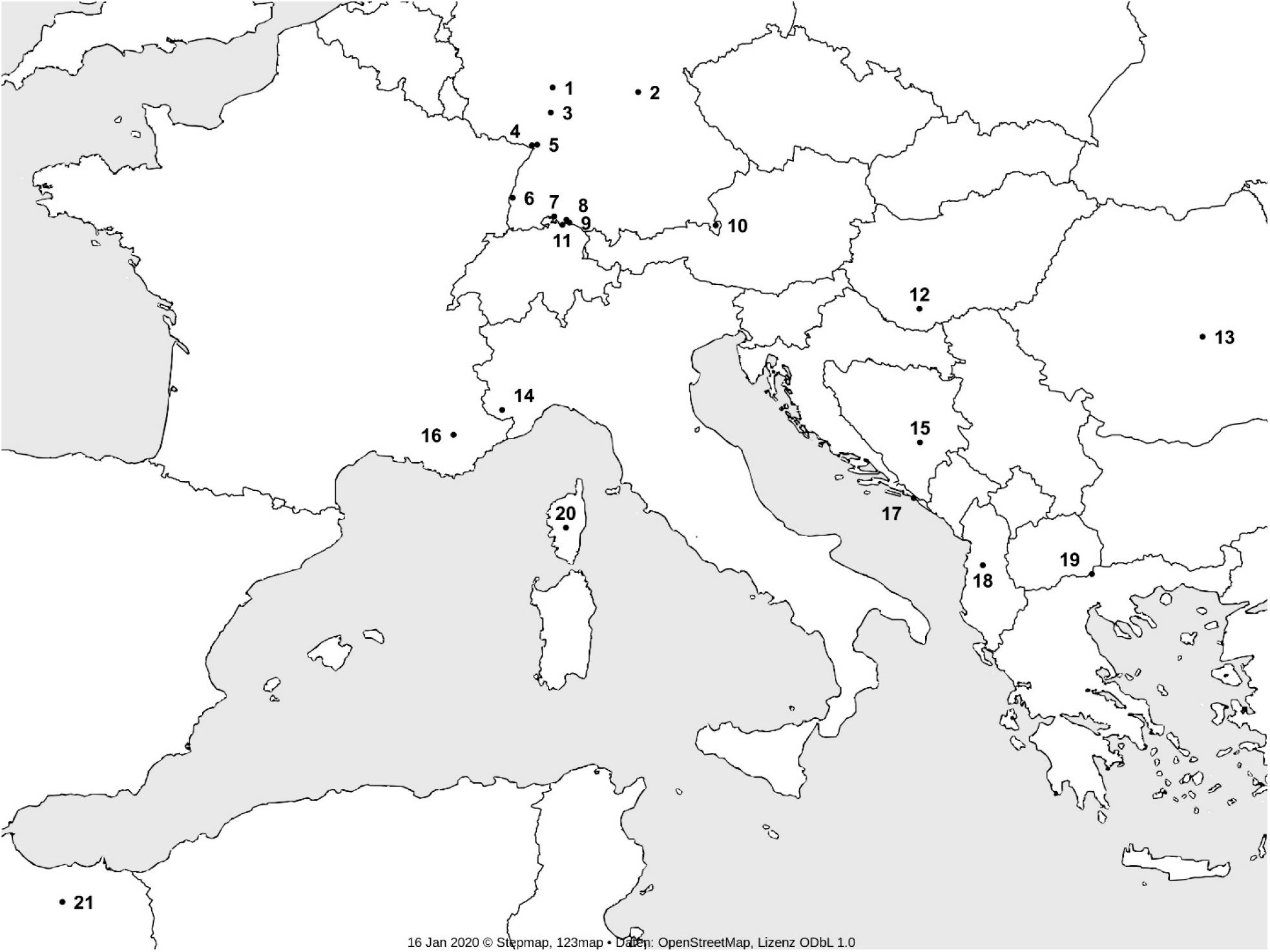
Origin of *S. saxatilis* museum specimens used for chemical analysis. Numbers in parentheses indicate numbers of specimens sampled per location (MFNB: Museum für Naturkunde, Berlin, Germany; SMNK: Staatliches Museum für Naturkunde Karlsruhe, Germany; SDEI: Senckenberg Deutsches Entomologisches Institut, Müncheberg, Germany). Countries on labels were translated. 1: Germany, Odenwald, Dreieichenhain, September 15^th^ 1925, MFNB (1); 2: Germany, Bamberg, Hallstadt, Börstig, August 3^rd^ 1940, leg. Schneid, MFNB (1); 3: Germany, Baden, Bergstraße, Weinheim, May 1952, leg. H. Nowotny, SMNK (1); 4: Germany, Bienwald, Büchelberg, September 10^th^ 1987, leg. Roesler, SMNK (2); 5: Germany, Karlsruhe, Maxau, August 22^nd^ 1959, leg. Kormann; Germany, Karlsruhe, Maxau, August 1947, leg. Nowotny, SMNK (2); 6: Germany, Baden, Kaiserstuhl, June 21^st^ 1953, leg. H. Nowotny, SMNK (1); 7: Germany, Baden, Hegau, August 8^th^-20^th^ 1935, leg. Leininger, SMNK (2); 8: Germany, Bodensee, Allensbach, September 1919, leg. W. Ramme, MFNB (2); 9: Germany, Wollmatingen, August 10^th^ 1928, leg. Leininger, SMNK (2); 10: Germany, Berchtesgaden, Bischofswiesen, August 21^st^-28^th^ 1958, leg. Papperitz, SDEI (1); 11: Switzerland, Glarisegg, May 6^th^-10^th^ 1906, SDEI (1); 12: Hungary, Mecsek-Gebirge, Mánfa, May 30^th^ 1977, leg. U. Göllner, MFNB (1); 13: Romania, Braşov, August 13^th^ 1905, leg. E.J. Lehmann, MFNB (1); 14: Italy, Alpi Marittime Natural Park, Juniperus phoenicea Riserva Naturale, August 10^th^ 2013, leg. J. Deckert, MFNB (2); 15: Bosnia and Herzegovina, Bjelašnica, July 21^st^ and 23^rd^ 1909, leg. F. Schumacher, MFNB (2); 16: France, Riez, July 1985, leg. J. Haupt, MFNB (1); 17: Croatia, Dubrovnik, April 13^th^ and May 1^st^ 1938, leg. Dr. Feige, SDEI (3); 18: Albania, Dajti Südhang, June 30^th^ 1961, leg. Expedition DEI, SDEI (1); 19: Republic of Macedonia, Hügel bei Stari Dojran, August 3^rd^ 1975, leg. U. Göllner, MFNB (1); 20: France, Corsica, Bocognano, 1905, leg. O. Leonhard, SDEI (1); 21: Kingdom of Morocco, Taza, MFNB (1).

**Supplemental Figure 6.**
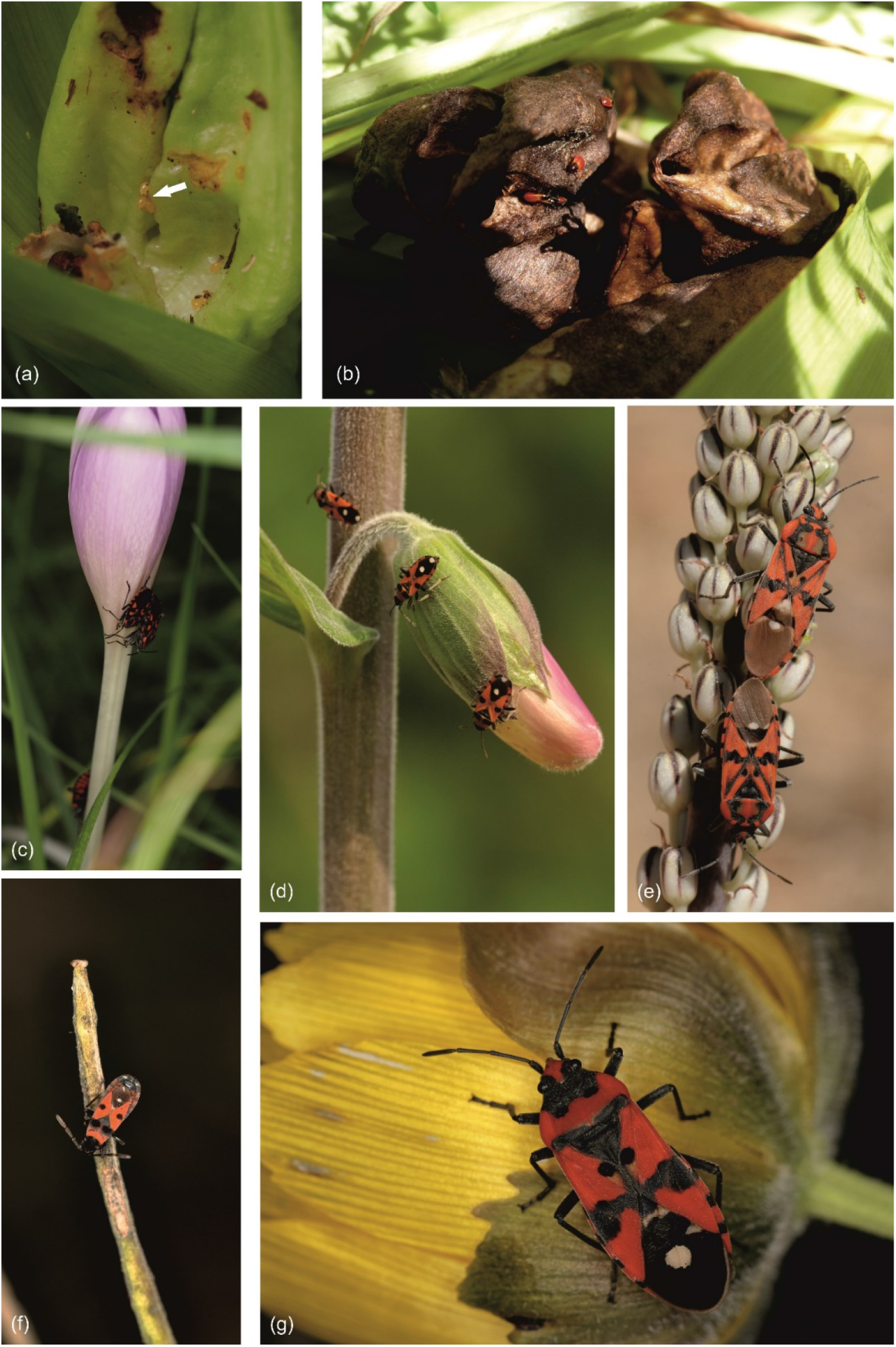
Natural history observations on milkweed bug species. (a) Eggs of *S. saxatilis* (white arrow) in a *C. autumnale* infructescence (Germany, Baden-Württemberg, Berghausen, June 1^st^, 2016). (b) Early instar larvae of *S. saxatilis* sitting on a dry *Colchicum* seedpod (Germany, Baden-Württemberg, Berghausen, June 15^th^, 2016). (c) Adults of *S. saxatilis* feeding on a *C. autumnale* flower (Germany, Baden-Württemberg, Berghausen, September 18^th^, 2015). (d) *H. superbus* on *D. purpurea* (Germany, Baden-Württemberg, Eberbach, June 5^th^, 2016). (e) *S. pandurus* on *U. maritima* (Spain, Sierra de Aracena and Picos de Aroche Natural Park, Cañaveral de León, August 20^th^, 2014). (f) *H. superbus* on *E. crepidifolium* (Germany, Rheinland-Pfalz, Schloßböckelheim, June 13^th^, 2016). (g) *L. equestris* on *A. vernalis* (Germany, Brandenburg, Mallnow, April 19^th^, 2016.

**Supplemental Figure 7.**
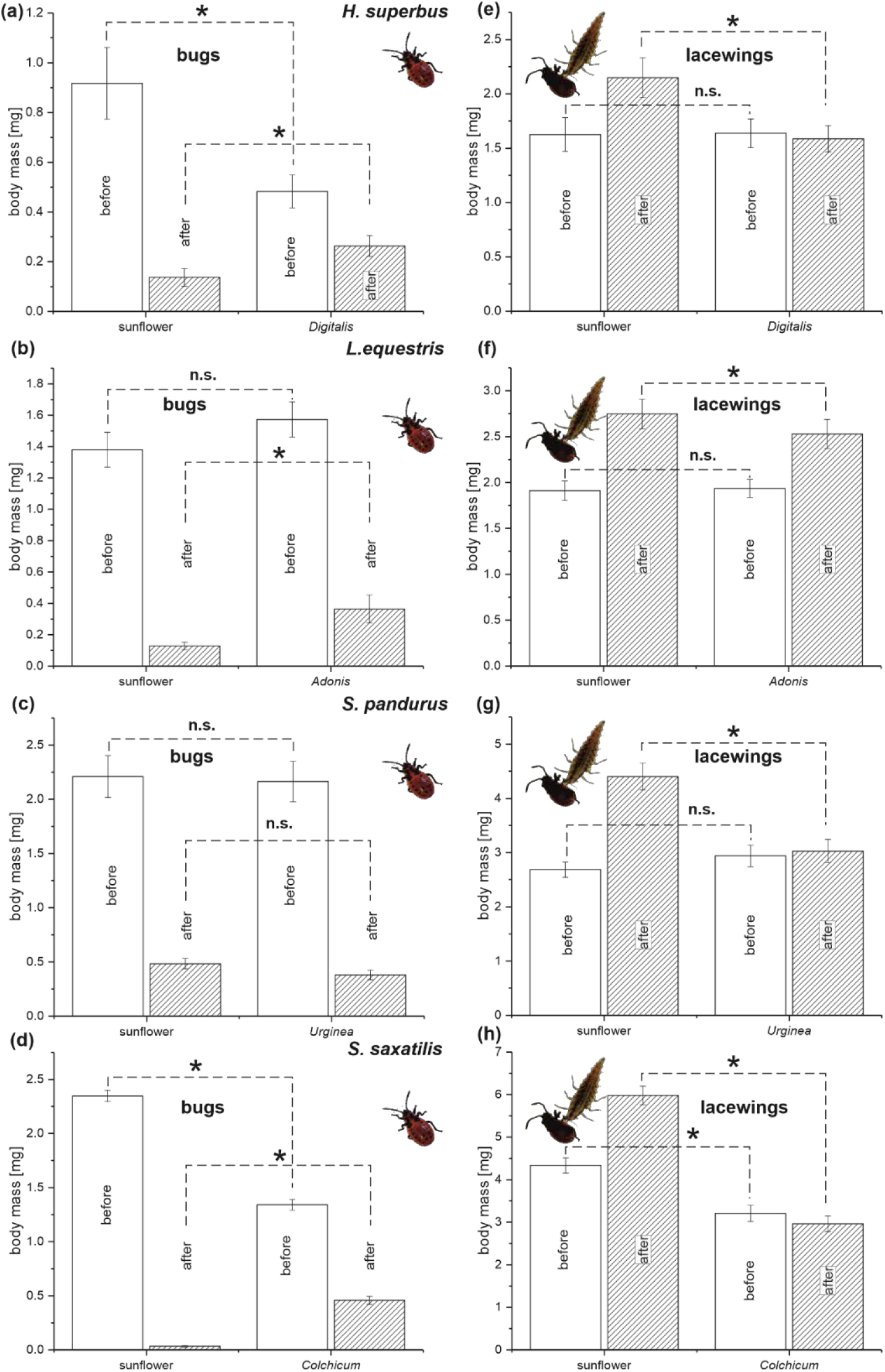
Consumption of four species of milkweed bugs by lacewing larvae (*C. carnea*). Early instar larvae of milkweed bugs were raised either on sunflower seeds (controls) or on seeds of plant species containing toxins for sequestration. Left panel: Body mass of milkweed bug larvae before (open bars) and after lacewing attacks (hatched bars). (a) *H. superbus* on *D. purpurea*, (b) *L. equestris* on *A. vernalis*, (c) *S. pandurus* on *U. maritima*, (d) S*. saxatilis* on *C. autumnale*. Right panel: Body mass of lacewing larvae before (open bars) and after feeding on milkweed bug larvae (hatched bars). (e) *H. superbus* on *D. purpurea*, (f) *L. equestris* on *A. vernalis*, (g) *S. pandurus* on *U. maritima*, (h) S*. saxatilis* on *C. autumnale*. Shown are means ± SE. Asterisks indicate significant differences at p = 0.05, n.s. = not significant.

**Supplemental Figure 8.**
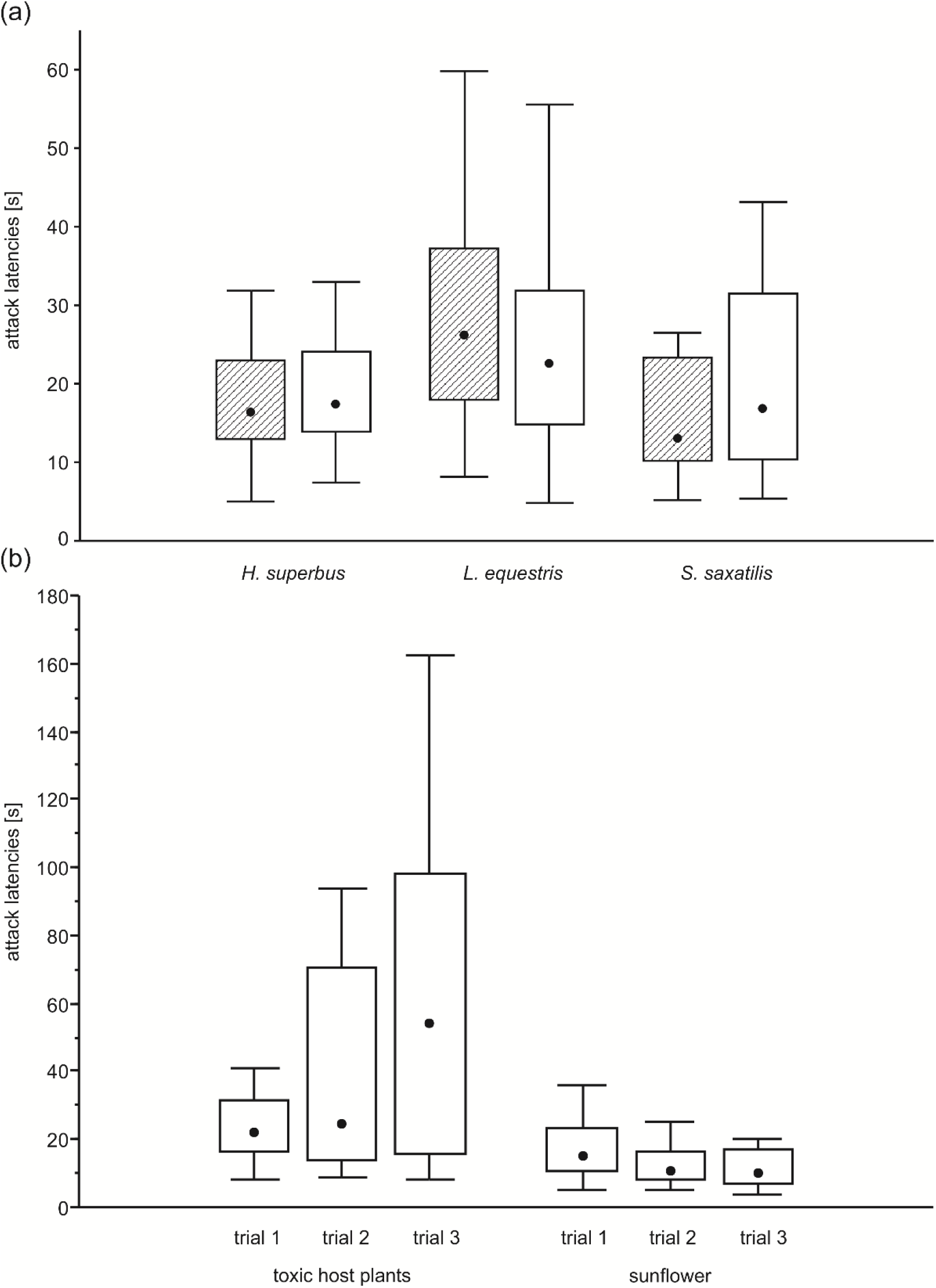
Attack latencies of juvenile great tits (*Parus major*) preying on milkweed bugs and control palatable prey. (A) Attack latencies in first encounters with novel palatable prey (crickets; hatched rectangles) and milkweed bugs (*H. superbus*, *L. equestris*, and *S. saxatilis*; open rectangles). (B) Changes in attack latencies across three successive trials with milkweed bugs raised either on seeds from toxic host plants or sunflower seeds as a control (data from *H. superbus*, *L. equestris*, and *S. saxatilis* pooled). Only the data from bugs that were actually attacked by birds in the respective trials are included. Boxes represent median (points), quartiles (rectangles) and range (whiskers).

**Supplemental Figure 9.**
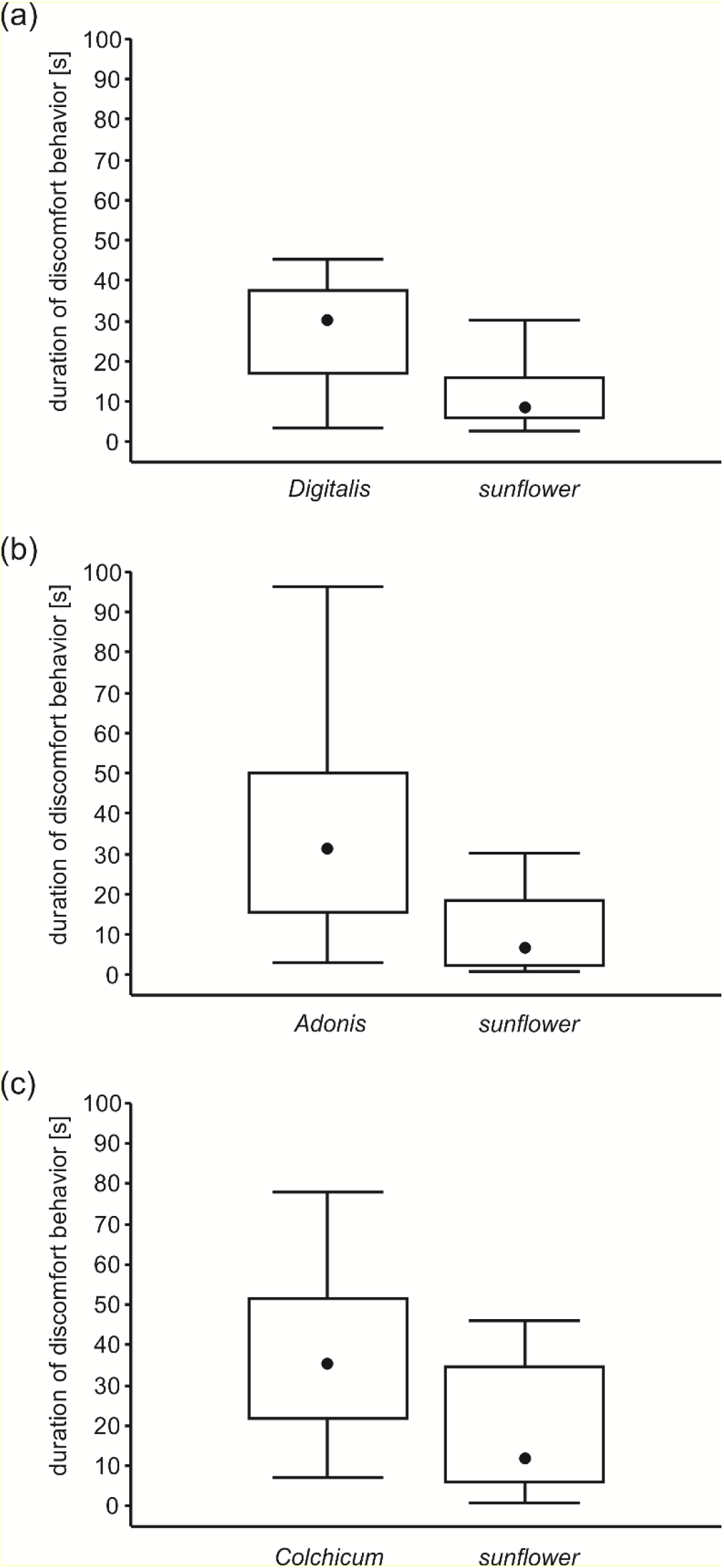
Durations of discomfort-indicating behaviors (beak wiping, head shaking) exhibited by juvenile great tits (*Parus major*) following their first contact (attack and handling) with bugs either raised on seeds of toxic host plants or sunflower as a control. a, b, c: birds tested with *H. superbus*, *L. equestris*, and *S. saxatilis*. Boxes represent median (points), quartiles (rectangles) and range (whiskers).

**Supplemental Figure 10.**
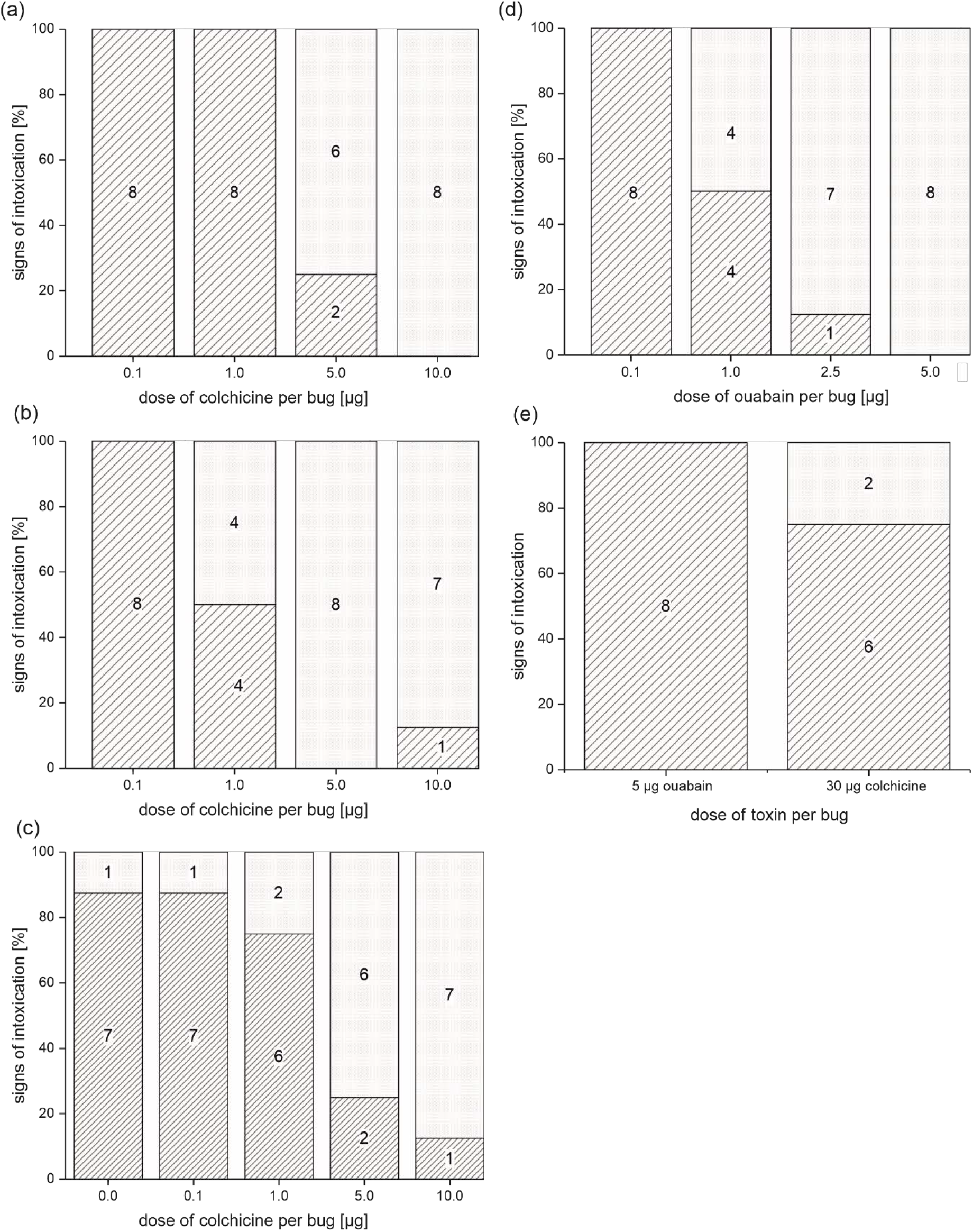
Resistance of *P. apterus, O. fasciatus, S. saxatilis*, and *S. pandurus* to injected ouabain and colchicine. (a) Effect of increasing doses of injected colchicine and ouabain (d) on *P. apterus*, (b) Effect of increasing doses of injected colchicine on *O. fasciatus*, (c) Effect of increasing doses of injected colchicine on *S. pandurus*. (e) Effect of injected ouabain and a high dose of colchicine on *S. saxatilis*. Bars show proportions of individuals that either showed signs of intoxication (open) or showed no signs of intoxication (hatched). Numbers in stacked bars represent the actual number of specimens.

